# Influenza A virus activates the unfolded protein response and induces the accumulation of insoluble protein aggregates that are essential for efficient viral propagation

**DOI:** 10.1101/2023.09.11.557148

**Authors:** Mariana Marques, Bruno Ramos, Hélio Albuquerque, Marisa Pereira, Diana Roberta Ribeiro, Alexandre Nunes, Jéssica Sarabando, Daniela Brás, Ana Rita Ferreira, Rui Vitorino, Maria João Amorim, Artur M. S. Silva, Ana Raquel Soares, Daniela Ribeiro

## Abstract

Influenza A virus (IAV) is one of the main causes of annual respiratory epidemics in humans. IAV employs multiple strategies to evade host immunity and hijack cellular mechanisms to support proper virion formation and propagation. Some of these strategies encompass the manipulation of pathways involved in protein homeostasis, leading to changes in the host proteome and protein distribution within the cell. In this study, we performed a detailed analysis of the interplay between IAV and the host cells’ proteostasis mechanisms throughout the entire infectious cycle. We reveal that IAV infection induces the activation of the inositol requiring enzyme 1 (IRE1) branch of the unfolded protein response (UPR), at an infection stage that coincides with high rates of viral protein translation. This activation is particularly important for infection, as attenuation of virus production was observed upon IRE1 inhibition. Concomitantly to UPR activation, we observed the accumulation of virus-induced insoluble protein aggregates, which contain both viral and host proteins and are associated with a dysregulation of the host cell RNA metabolism. We demonstrate that this accumulation is important for IAV propagation, as its prevention using a quinoline-steroid hybrid compound significantly reduces the number of produced infectious virus particles. Our data suggests that the formation of these insoluble protein aggregates favors the final steps of the infection cycle, more specifically the virion assembly. Our findings reveal additional mechanisms by which IAV disrupts the host cell proteostasis to favor infection and uncover new cellular targets that can be explored for the development of host-directed antiviral strategies.

## Introduction

Viruses have developed multiple strategies to hijack and control host cellular activities to evade the immune response and support efficient virus particle production. Some of these strategies include the manipulation of cytoplasmic and endoplasmic reticulum (ER)-related mechanisms involved in protein metabolism [1–3]. To counteract disturbances in the proteome and re-establish basal homeostasis, cells have evolved distinct surveillance mechanisms concerning protein biogenesis, folding, degradation, and sequestration of abnormal and potentially pathogenic conformers [4]. One of the most important mechanisms for protein stress detection is the ER unfolded protein response (UPR) [5]. Upon activation of one or more of the key UPR signal activator proteins (inositol requiring enzyme (IRE1), protein kinase R (PKR)-like ER kinase (PERK) and activating transcription factor 6 (ATF6)), downstream signaling leads to the attenuation of general protein synthesis with selective protein translation (via PERK), preferential degradation of mRNA encoding for ER-localized proteins (via IRE1), and the controlled synthesis of stress-attenuating proteins, such as chaperones and folding catalysts (via PERK, IRE1 and ATF6) [6]. The ER-associated protein degradation (ERAD) pathway may also be activated, allowing the clearance of misfolded and accumulated proteins in the ER [7]. Various cytoplasmic chaperones, such as heat shock proteins (HSPs), also assist the folding and assembly of newly synthesized and stress-damaged proteins, to prevent potentially pathogenic protein aggregation [8]. Misfolded or aggregated proteins may also be sequestered and compartmentalized into stress foci to hinder their toxic effects and minimize interference [9].

Several viruses lead to the accumulation of protein aggregates in specialized compartments, generally termed virus factories, in order to recruit and concentrate viral and host components and facilitate the molecular interactions required for essential steps of genome replication or virus particle assembly, while escaping immune recognition [10]. As part of the host antiviral response to infection, cytoplasmic aggregates, comprising both host and viral components, can also be formed and eventually become targeted for degradation to facilitate viral clearance and cellular recovery [11].

The importance of protein homeostasis in the context of viral infections is well recognized, supporting that further efforts to understand the underpinning molecular mechanisms may lead to antiviral control. With that in mind, in this work we performed a detailed analysis of different proteostasis-related mechanisms in the course of influenza A virus (IAV) infection. IAV is the causative agent of most of the annual respiratory epidemics in humans [12], with a high level of morbidity and mortality in the elderly and individuals with chronical disease conditions [13]. IAV belongs to the *Orthomyxoviridae* family, featuring a segmented genome composed by eight single-stranded negative-sense linear RNA segments, separately enclosed and wrapped by the viral nucleoprotein (NP) in the form of viral ribonucleoprotein complexes (vRNPs) [14]. Upon successful binding to the host cell membrane, IAV virions are internalized and its genome is released into the cytoplasm, being further imported to the nucleus where it undergoes transcription and replication [15]. The translation of viral proteins at the cytoplasm and the ER, together with the formation of vRNPs in the nucleus, culminate in the assembly of progeny virus that bud at the plasma membrane before being released from the cell [15].

Not much is known concerning the interplay between IAV and the host cell proteostasis, and the available information is somewhat contradictory. Regarding the UPR, while some authors reported the activation of the ATF6 pathway upon IAV infection [16] and others suggested the inhibition of PERK as a possible IAV antiviral strategy [17], additional studies have described the stimulation of the IRE1 branch upon infection [18–21] with little or no concomitant activation of the ATF6 and PERK branches [20,21]. These variances may, however, be related to the use of different cell models and/or viral strains. Nevertheless, the UPR has been suggested as a putative target for host-directed antivirals against IAV infection, as its activation by thapsigargin has been shown to block viral replication [22]. More recently, UPR activation mediated by thiopurines led to the selective disruption of the synthesis and maturation of IAV glycoproteins HA and NA, eventually blocking viral replication [23]. Some studies have described the accumulation of IAV-derived amyloid-like fibers to induce cytotoxicity [24], as well as the formation of aggresome-like structures in IAV-infected dendritic cells to evade the immune response [25]. It has also been shown that the accumulation of cytosolic stress granules, normally formed upon different kinds of cellular stress, is prevented upon IAV infection [26]. To clarify how the host cell proteostasis impacts IAV infection, we evaluated how specific proteostasis-related mechanisms are affected throughout the main phases of a single IAV infectious cycle. Our results not only demonstrate that IAV interferes with the UPR at different stages of infection, mainly through its IRE1 branch, but also that this interplay is required for proper virus particle formation and propagation. Importantly, we demonstrate that, upon high rates of viral protein translation, IAV induces the accumulation of insoluble protein aggregates at the cytosol composed of both host and viral proteins. By chemically disrupting the assembly of these protein aggregates, we demonstrate that this process is essential for efficient viral protein production and proper formation of infectious virus particles, supporting the idea that targeting proteostasis-related mechanisms may constitute a valid therapeutic approach to tackle and decrease viral propagation.

## Results

### Influenza A virus triggers the IRE1 branch of the unfolded protein response to ensure efficient viral propagation

To elucidate which UPR pathways are specifically activated or remodeled throughout the different steps of the virus infectious cycle, we performed a detailed analysis of the distinct UPR signaling pathways (depicted in **Fig. 1A**) at different time points during a single cycle of infection. This comparative approach allowed assessing the interplay between infection and host proteostasis mechanisms and its potential to be used as a therapeutic target.

**Figure 1.**
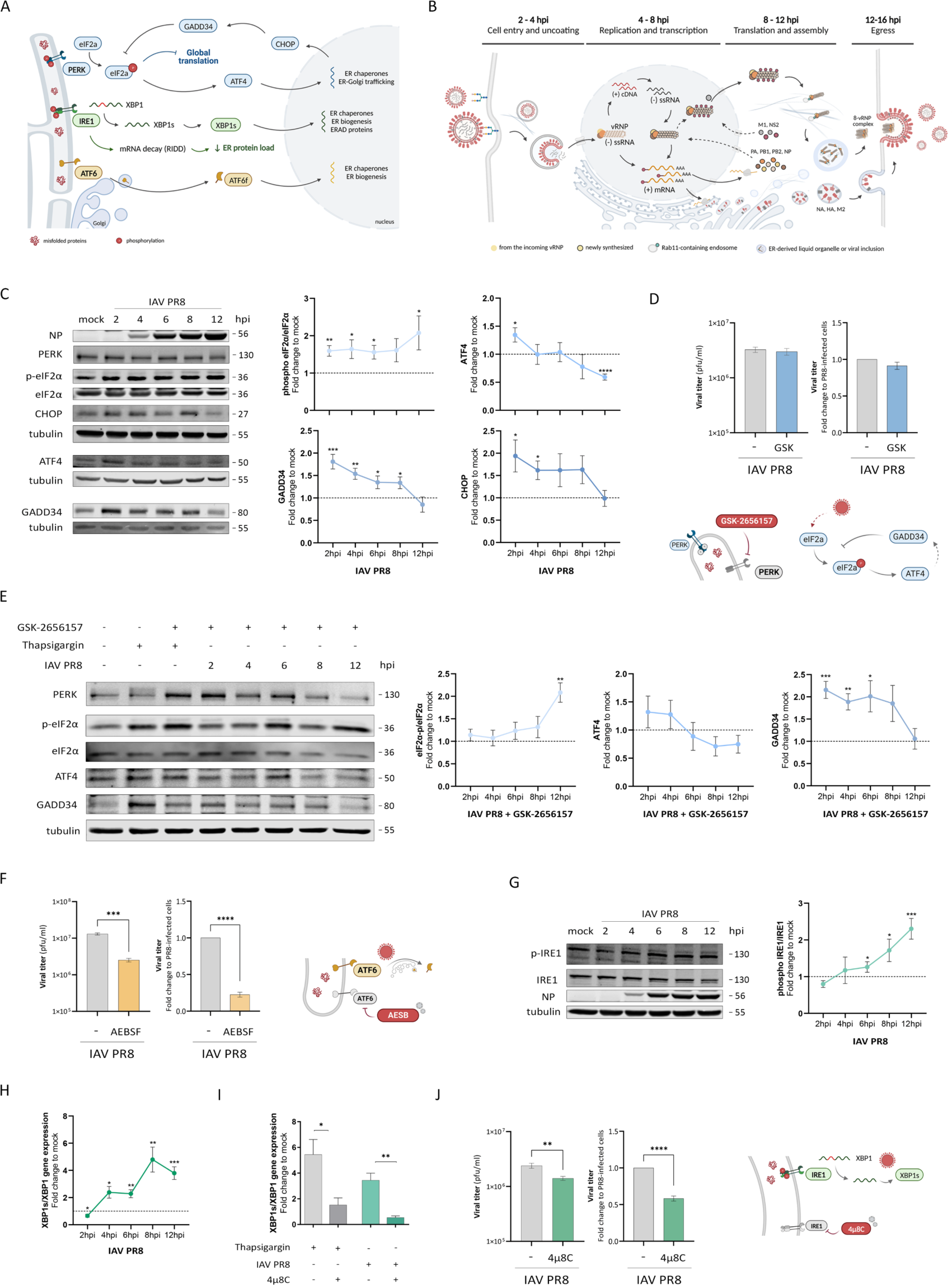
| Analysis of the host cell UPR during IAV infection. (A) Schematic representation of the three main branches of UPR in the ER. (B) Schematic representation of the influenza A virus life cycle. Important steps in infection are indicated with the usual time frames post infection in A549 cells. (C – E) Analysis of the relevance of the UPR PERK branch during infection with IAV in A549 cells. (C) Western blot analysis of the expression levels of PERK, p-eIF2α/eIF2α, CHOP, ATF4 and GADD34 proteins in A549 cells, following IAV PR8 infection at different times post infection. Tubulin was used as internal control. (D) Similar analyses as in C, but in the presence of 0.33 µM of the PERK inhibitor GSK-2656157. 500 nM thapsigargin treatment for 8 hours served as positive control for PERK activation. Quantification values were normalized to mock and represent average ± SEM of at least three independent experiments. Tubulin was used as internal control. (E) Plaque assay analysis of the viral titer obtained upon IAV PR8 infection of A549, in the presence or absence of 0.33 µM GSK-2656157. Data represents the means ± SEM of three independent experiments. (F) Analysis of the relevance of the UPR ATF6 branch during infection with IAV in A549 cells. Plaque assay analysis of the viral titer of A549 cells infected with IAV PR8 in the presence or absence of AEBSF (ATF6 inhibitor). (G – J) Analysis of the relevance of the UPR IRE1 branch during infection with IAV in A549 cells. (G) Western blot analysis of the expression levels of the phosphorylated and total forms of the IRE1 protein at different times post infection. Quantification values were normalized to mock-infected cells and tubulin was used as internal control. (H) RT-qPCR analysis of the splicing of XBP1 at different times post infection in relation to mock-infected cells and (I) upon stimulation with thapsigargin or infected with IAV PR8, in the presence or absence of 4μ8C (IRE1 inhibitor), in relation to untreated cells. (J) Plaque assay analysis of the viral titer obtained upon IAV PR8 in the presence or absence of 4μ8C. Data represents the means ± SEM of at least three independent experiments. *p<0.005, **p<0.001, ***p<0.0001, ****p<0.00001 using student’s t-test.

Adenocarcinomic human alveolar basal epithelial (A549) cells were infected with influenza A/Puerto Rico/34/8 (IAV PR8) [27,28] and UPR-related genes or proteins were assessed at different time-points post infection, reflecting the most relevant steps of the virus life cycle (depicted in **Fig. 1B**), namely 2, 4 6, 8 and 12 hpi. To investigate the PERK UPR branch, we evaluated the expression of several proteins belonging to this pathway, namely eIF2α and its phosphorylation, ATF4, CHOP and GADD34. eIF2α, which catalyzes an early step of protein synthesis initiation, becomes phosphorylated upon PERK activation to induce global translation arrest, while allowing the selective translation of different proteins such as ATF4 [5]. ATF4 further induces both CHOP and GADD34 expression to ultimately counteract eIF2α phosphorylation in a negative loop to restore translation levels (**Fig. 1A**). Quantification of the expression ratio between the phosphorylated and unphosphorylated eIF2α (**Fig. 1C**) showed an increase in eIF2α phosphorylation already at 2 hpi, being this activation maintained throughout the remaining infection cycle, as reported by others [29]. ATF4 expression increases at the initial steps of infection and decreases thereafter (**Fig. 1C**). Similarly, the expression of both CHOP and GADD34 increases at early time points and subsequently decreases progressively until the end of infection (**Fig. 1C**), although maintaining a higher level than in uninfected (mock) cells. However, the characteristic band smear that indicates PERK activation is not detected by western blotting upon infection (**Fig. 1C**).

To further elucidate the importance of the PERK pathway for IAV infection, cells were infected with IAV PR8 in the presence of a specific inhibitor of PERK, GSK-2656157 (as in [30]). Upon assessment of the amount of infectious virus particles produced in the presence and absence of this inhibitor, our results clearly indicate that PERK inhibition does not affect virus production (**Fig. 1D**). In addition, upon IAV infection, eIF2α phosphorylation is still observed upon incubation with the PERK inhibitor, and the expression levels of ATF4 and GADD34 are also maintained (**Fig. 1C and E**). Overall, this indicates that the observed eIF2α phosphorylation upon IAV infection is unlikely to result from PERK activation, but rather from the activation of another kinase.

In order to analyze the relevance of the UPR ATF6 pathway for IAV infection, we infected the cells upon incubation with a broad-spectrum serine protease inhibitor commonly used to inhibit ATF6, 4-(2-aminoethyl) benzenesulfonyl fluoride (AEBSF) (as in [20,31]). Quantification of the formation of infectious virus particles by plaque assay demonstrated a significant decrease in the number of new virus particles upon inhibition of the ATF6 pathway (**Fig. 1F**). AEBSF prevents ER stress-induced cleavage of ATF6α and ATF6β, resulting in inhibition of transcriptional induction of ATF6-target genes [32], supporting the assumption that the ATF6 pathway plays an important role in IAV infection.

Lastly, the activation of the IRE1 pathway was assessed via the phosphorylation of IRE1 and the splicing of XBP1. We observed an increase in the phosphorylation of IRE1 starting at 6 hpi up to 12 hpi with IAV PR8 (**Fig. 1G**), reflecting a stage of the life cycle where viral proteins are actively being translated. Coherently, the levels of spliced XBP1 were significantly increased, (**Fig. 1H**), particularly at 8 hpi. To further infer the importance of this activation for viral propagation, we have quantified infectious virus particle formation in the presence of a specific inhibitor of this pathway, 4µ8C (as in [20,33]) (**Fig. 1I and J**). In the presence of this inhibitor, virus production was attenuated by approximately 40% (**Fig. 1J**), indicating that the activation of the UPR IRE1 branch by IAV is also relevant for IAV particle formation and propagation.

Collectively, our data indicates that UPR activation occurs at different stages of infection, and that efficient ATF6 and IRE1 pathways are required for the proper disenrollment of the IAV PR8 infection cycle.

### Influenza A virus induces the accumulation of cytosolic protein aggregates

We proceeded our study by analyzing whether IAV infection leads to proteostasis imbalances through the accumulation of insoluble proteins. A549 cells were infected with IAV PR8 and samples were harvested to further analyze the detergent-insoluble protein fractions by SDS-PAGE, as previously reported [34]. Insoluble proteins accumulate in cells infected with IAV (**Fig. 2A**), particularly at 8 hpi when a considerable amount of the viral RNA has been transcribed and replicated, viral transcripts and viral progeny RNA (in the form of vRNPs) have exited the nucleus, and a substantial amount of viral proteins is being synthesized. Our results indicate proteostasis impairment and overlap with the activation of the IRE1 pathway. This effect appears to be reversible at later times post-infection (**Fig. 2A**).

**Figure 2.**
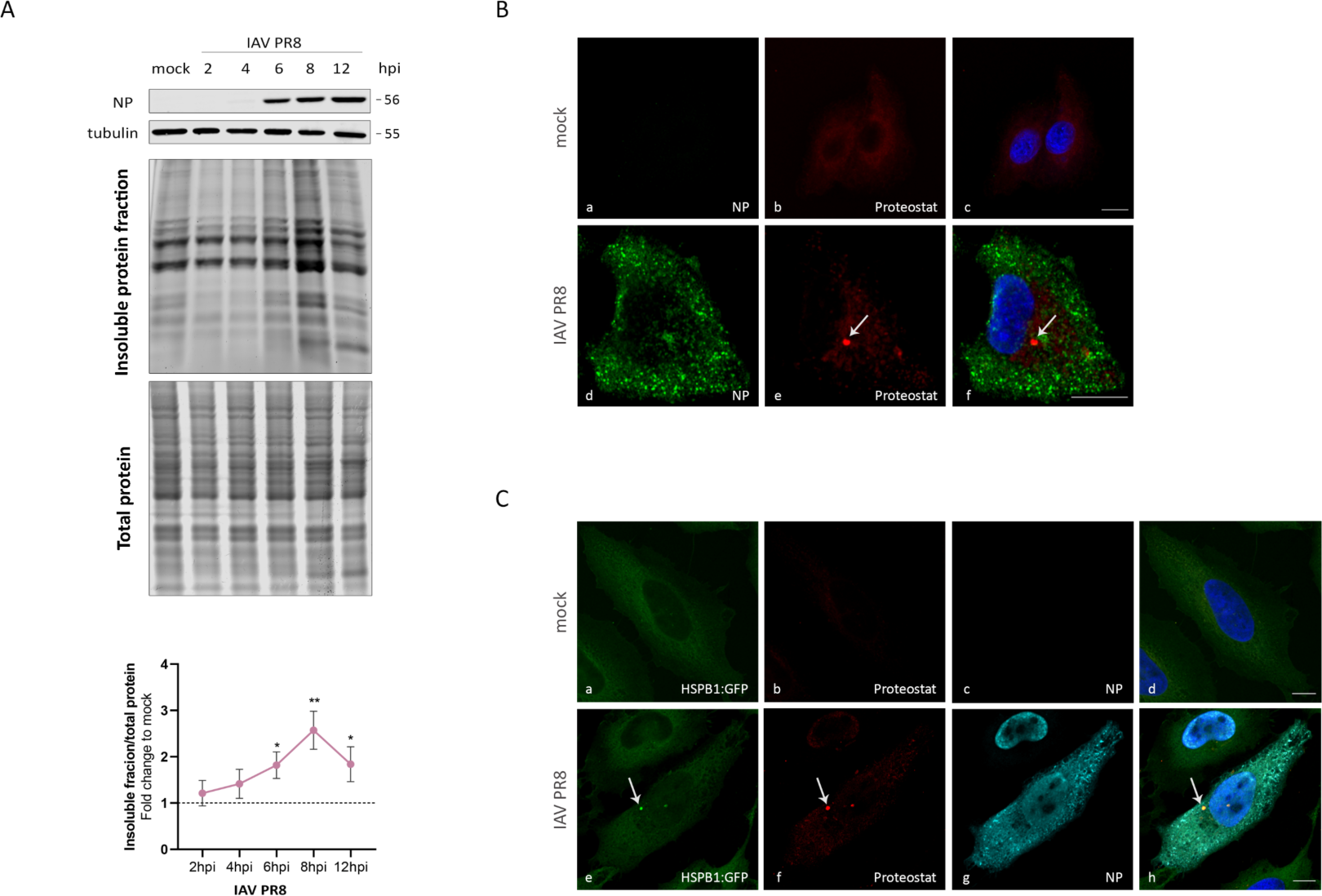
| Analysis of the accumulation of misfolded proteins in IAV-infected cells. (A) Characterization of the insoluble protein fraction of A549 cells infected with IAV PR8 at different times post infection. Data represents means ± SEM of at least independent experiments, *p<0.05, ****p<0.0001 using student t-test. (B) Aggresomes formation in mock- or IAV PR8-infected A549 cells at 8 hpi. Confocal images of (a, d) viral NP, (b, e) Proteostat® dye and (c, f) merge image. (C) Aggresomes formation in mock- or IAV PR8-infected in HeLa HSPB1:GFP cells at 8 hpi. Confocal images of (a, e) endogenous HSPB1:GFP, (b, f) Proteostat® dye, (c, g) anti-NP and (d, h) merge images. Arrows indicate the presence of aggresome-like structures. Bars represent 10 µm.

Aiming at visualizing and intracellularly localizing these protein aggregates, we stained A549 infected cells with Proteostat®, a dye that specifically marks for misfolded and aggregated proteins (**Fig. 2B**). At 8 hpi, Proteostat® stains several cytoplasmic protein aggregates (**Fig. 2B**), mainly localized at the perinuclear area. To solidify these results, we have performed the same experiment in another cell type, a previously established HeLa cell line that stably expresses a GFP-tagged protein misfolding sensor HSP27:GFP (HeLa HSP27:GFP cells) [35]. Similarly to A549 cells, the formation of several cytoplasmic protein aggregates stained by both Proteostat® and HSP27:GFP was observed (**Fig. 2C**).

Altogether, these results reveal that IAV induces the formation of cytoplasmic protein aggregates, at a time point of the infection cycle where an intense production of viral proteins is occurring [36], as well as the activation of the UPR IRE1 pathway.

### The insoluble protein fraction of infected cells contains viral proteins and is enriched in host translation-related proteins

To obtain further insights on the composition of these protein aggregates, we performed liquid chromatography tandem mass spectrometry analysis (LC-MS/MS) of the detergent-insoluble protein fraction of mock and IAV PR8-infected cells at 8 hpi (**Fig. 3A**).

**Figure 3.**
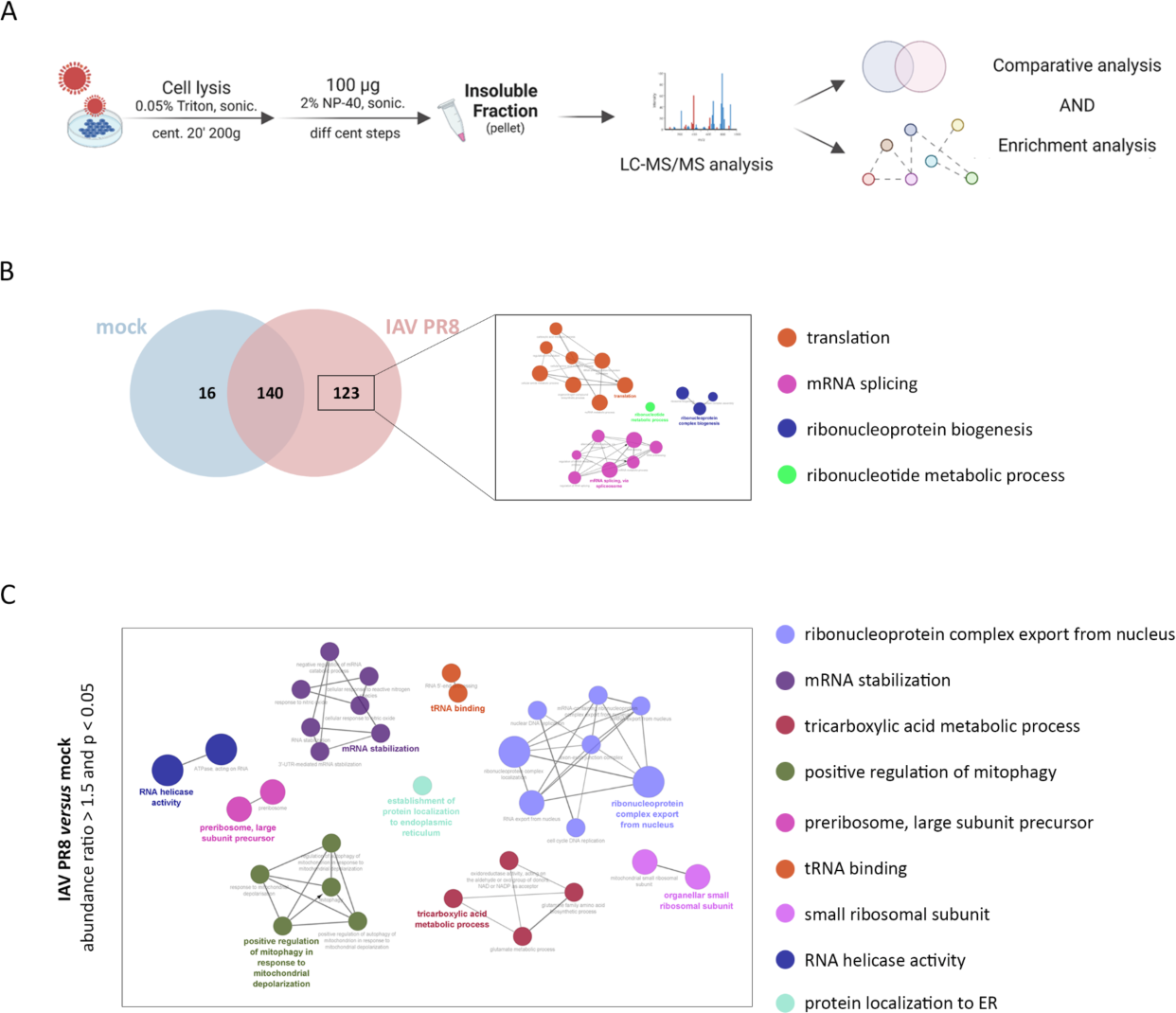
| Analysis of the host proteins present in the insoluble fraction of infected cells by LC-MS/MS. (A) Experimental approach used to isolate and characterize the insoluble protein fraction. (B-C) Comparison between the insoluble protein fractions in IAV PR8- and mock-infected cells. (B) Venn diagram representing the number of host’s proteins identified in the insoluble fractions from IAV PR8-infected and non-infected cells. Characterization of insoluble proteins found solely in the insoluble fraction of IAV PR8-infected cells using the ClueGo plugin in Cytoscape, based on the results from the LC-MS/MS analysis of three independent experiments. For this analysis, proteins identified by peptides and unique peptides >2 in at least 2 experiments were considered. (C) Gene ontology analysis (using ClueGO) of the enriched insoluble proteins in IAV PR8-infected cells.

Five viral proteins were found in the insoluble protein fraction of infected cells, namely the RNA-directed RNA polymerase catalytic subunit (PB1), matrix protein 1 (M1), nucleoprotein (NP), non-structural protein 1 (NS1) and polymerase basic protein 2 (PB2) (Supplementary Table 1). A total of 156 and 263 human proteins were identified with at least 2 unique peptides in mock and IAV PR8-infected cells, respectively (Supplementary Table 2), corroborating our previous results that show an increase in the level of insoluble proteins at this stage of infection. Of the identified proteins, 123 were solely found in the insoluble fraction of cells infected with IAV PR8 (**Fig. 3B**). A gene ontology analysis using the Cytoscape plug-in ClueGo revealed that these proteins belong to a specific set of biological processes related to translation, mRNA splicing and the ribonucleoprotein complex biogenesis (**Fig. 3B**).

Through a complementary analysis, we determined which host insoluble proteins were enriched in IAV PR8-infected cells in comparison to non-infected cells. Proteins with an abundance ratio (IAV PR8 to mock) greater than 1.5 with an adjusted *p*-value smaller than 0.05 were considered enriched. This analysis allowed the identification of 78 host proteins whose abundance is increased in the insoluble protein fraction upon infection (Supplementary Table 3). Gene ontology analysis using Cytoscape showed that these proteins are involved essentially in processes related to ribonucleoprotein complex export from the nucleus, mRNA stabilization, but also in processes such as the regulation of mitophagy, or the tricarboxylic acid metabolism (**Fig. 3C** and Supplementary Table 4A). An extra analysis using STRING database showed that our network of proteins enriched in the insoluble fraction of IAV PR8-infected cells are related to nuclear DNA replication, ribonucleoprotein export from the nucleus and several processes related to RNA metabolism (Supplementary Table 4B).

These results indicate that the solubility of several host proteins related to protein translation and RNA processing is altered at this time point of infection. Whether this occurs to benefit viral infection or as a host strategy to counteract infection is still unclear.

Viral and host proteins found to be enriched in the insoluble fraction of IAV PR8-infected samples in comparison to mock (Supplementary Tables 1 and 3) were then accessed for their natural propensity to aggregate, using PASTA 2.0 [37], a web server for the prediction of protein aggregation propensity by determining fibril formation. Beta-amyloid and bovine serum albumin (BSA) were used respectively as positive and negative controls of aggregation propensity and to define the threshold of best energy values for further comparison analyses. As depicted in **Table 1**, all IAV proteins identified in the insoluble fractions of infected cells have no propensity to aggregate, namely PB1, M1, NP, NS1 and PB2. On the other hand, only 10 of the 78 proteins from the host have shown to have propensity to aggregate (defined as having best energy < - 11), which corresponds to 12.8% of the detected host proteins in this fraction. These results indicate that most of the detected proteins in the insoluble fractions of infected cells are indeed accumulating as a result of the viral infection.

**Table 1.**
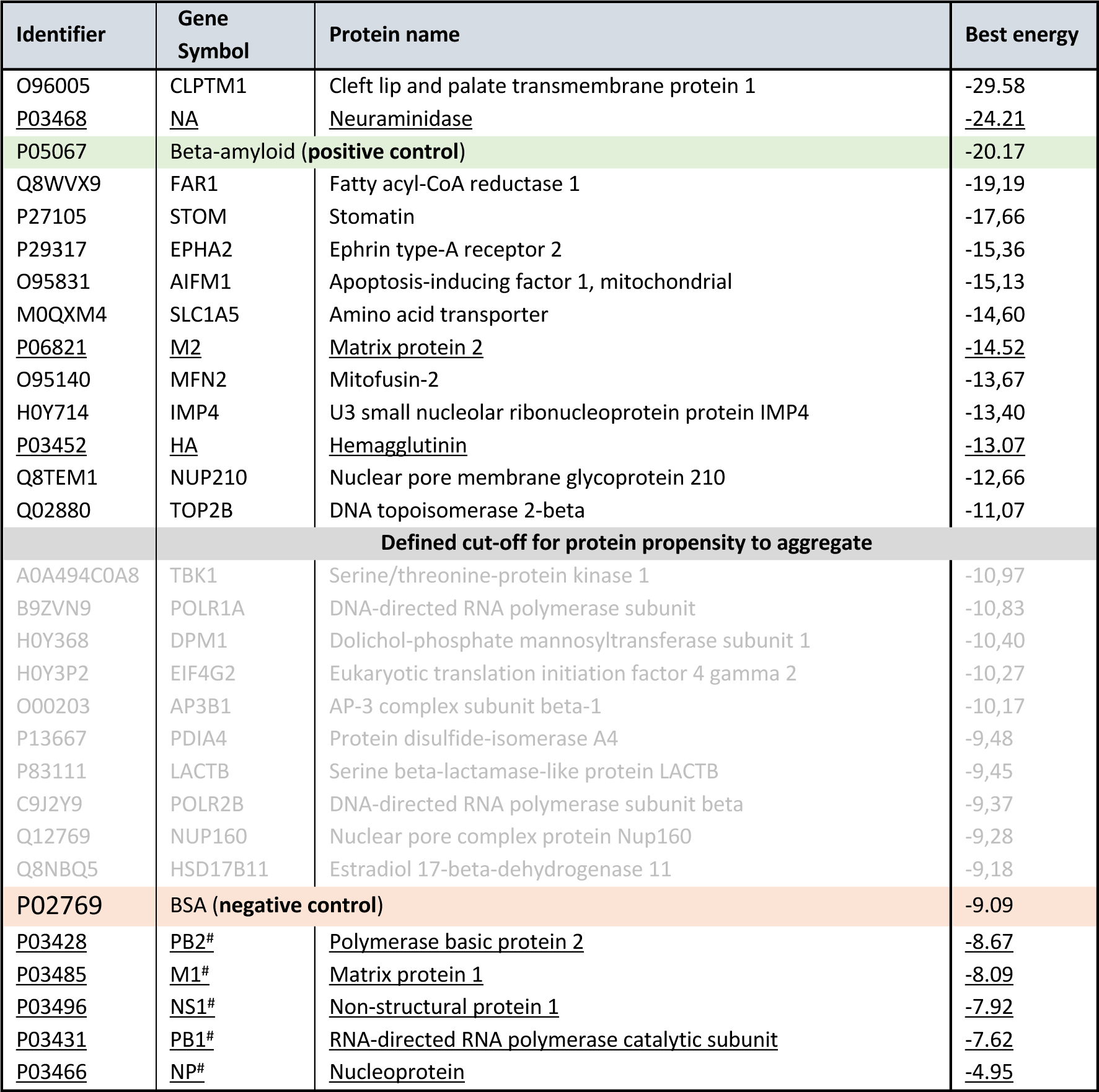
| Aggregation prediction of host’s and influenza A virus’ (underlined) proteins identified in the insoluble fractions of infected cells using PASTA 2.0. Beta-amyloid and BSA were used as a positive of negative control, respectively. Host proteins with best energy values higher than BSA but below cut-off are represented in grey; host proteins with best energy values below BSA are not represented. # refers to viral proteins found in the insoluble protein fraction of infected cells.

### The formation of IAV-induced protein aggregates favors infection

To understand if the observed virus-induced accumulation of insoluble protein aggregates is relevant for the process of virus particle formation, we disturbed the assembly of these aggregates and analyzed its consequence for viral propagation. For that, we used HASQ-6Cl (C_42_H_52_ClNO, ^1^H and ^13^C-NMR spectra (Supplementary Fig. 1), a steroid-quinoline hybrid that has been shown to disrupt and revert protein aggregation processes (named 6c in [38]) (**Fig. 4A**). This compound was tested for cell viability and toxicity and the ideal experimental concentration was determined (Supplementary Fig. 1).

**Figure 4.**
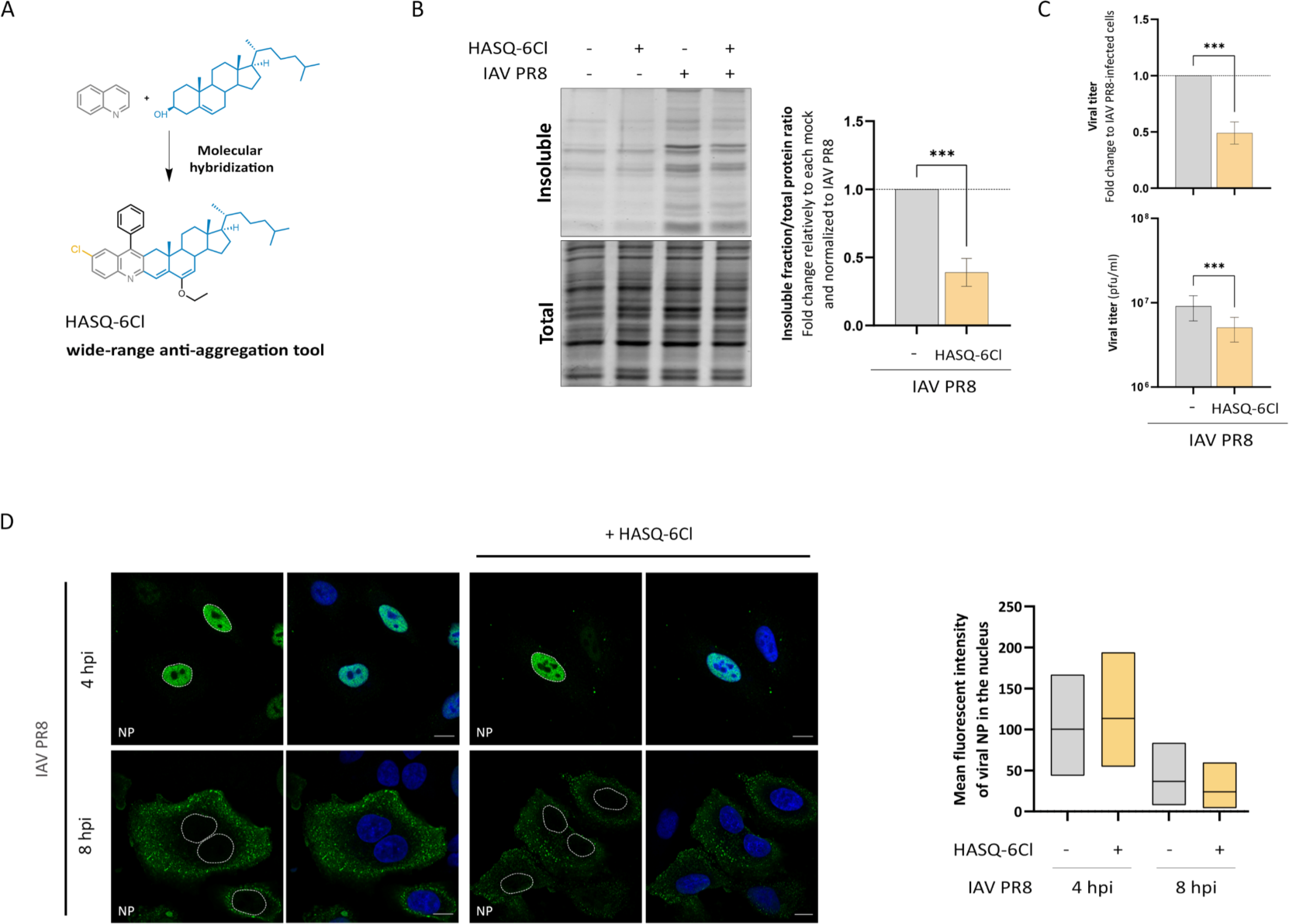
| Importance of protein aggregation for IAV propagation. (A) Simplified representation of the molecular hybridization reaction used to synthesize HASQ-6Cl. The combination of quinolines (in gray) and steroids (in blue) in one new chemical entity, generates a hybrid molecule with wide-range of anti-aggregation functions. (B) Analysis of the insoluble fraction of A549 cells infected for 8 h with IAV PR8 in the absence or presence of HASQ-6Cl. The final ratios were obtained by first normalizing the intensity of insoluble proteins fractions to the intensity of total fraction, followed by its normalization to the correspondent mock, and finally its normalization to IAV PR8. (C) Viral titers of IAV PR8 after treatment of A549 cells with HASQ-6Cl. Values were normalized to IAV PR8 and represent average ± SEM of at least three independent experiments, ***p<0.001, in student’s t-test. (D) Confocal images of A549 cells infected with IAV PR8 for 4 h and 8 h in the absence or in the presence of HASQ-6Cl. Viral NP is stained in green and nuclei are in blue (DAPI). Bars represent 10 µm. Data represents the mean fluorescence intensity of viral NP in the nucleus as box and whiskers min to max of three independent experiments, ****p<0.0001 using ordinary one-way ANOVA followed by Bonferroni’s multiple comparisons test.

A549 cells were treated with 50 µM of HASQ-6Cl 12 h prior to infection with IAV PR8 and harvested at 8 hpi (the time point after infection where we observed a higher accumulation of insoluble proteins) to analyze the insoluble protein fraction. The virus-induced levels of insoluble proteins decreased around 50% in the presence of HASQ-6Cl (**Fig. 4B**), demonstrating that this compound is also able to inhibit the formation of IAV-induced protein aggregates. The effect of this protein aggregation-inhibition on IAV infectious particle formation was analyzed by plaque assay at 16 hpi. A decrease of around 50% in the number of infectious viral particles in the presence of HASQ-6Cl was observed (**Fig. 4C**), indicating that the virus infection cycle is disturbed by the disaggregating compound.

To infer the specific infection step that was affected by the compound, we analyzed the viral NP mean fluorescence intensity (MFI) in the nucleus of IAV PR8-infected A549 cells, in the presence and absence of HASQ-6Cl. Our results indicate that the amount of nuclear NP is not affected by HASQ-6Cl at 4 or 8 hpi (**Fig. 4D**) and further show that HASQ-6Cl does not interfere either with the IAV entry into the cell or with the export of vRNPs from the nucleus at later times post-infection, suggesting a downstream effect on the viral life cycle.

Although these results cumulatively suggest that the formation of these protein aggregates may be a virus-induced mechanism to favor viral protein production, one should also hypothesize whether the decrease in virus titer in the presence of HASQ-6Cl could also result from its effect on a host mechanism that further impairs viral infection. As we had previously demonstrated that the activation of the UPR IRE1 branch is relevant for proper IAV particle formation, we questioned whether HASQ-6Cl would exert some effect over this pathway and consequently cause the observed decrease in virus titers. To investigate this hypothesis, we measured the splicing of XBP1 mRNA upon 8 h of infection with IAV PR8 in the presence or absence of HASQ-6Cl. **Fig. 5A** shows that there were no observed differences in XBP1 splicing in the presence of HASQ-6Cl, demonstrating that the compound has no effect on the level of IRE1 pathway induction by the virus. Our results also show no significant changes in the level of eIF2α phosphorylation throughout infection in the presence of HASQ-6Cl (**Fig. 5B**).

**Figure 5.**
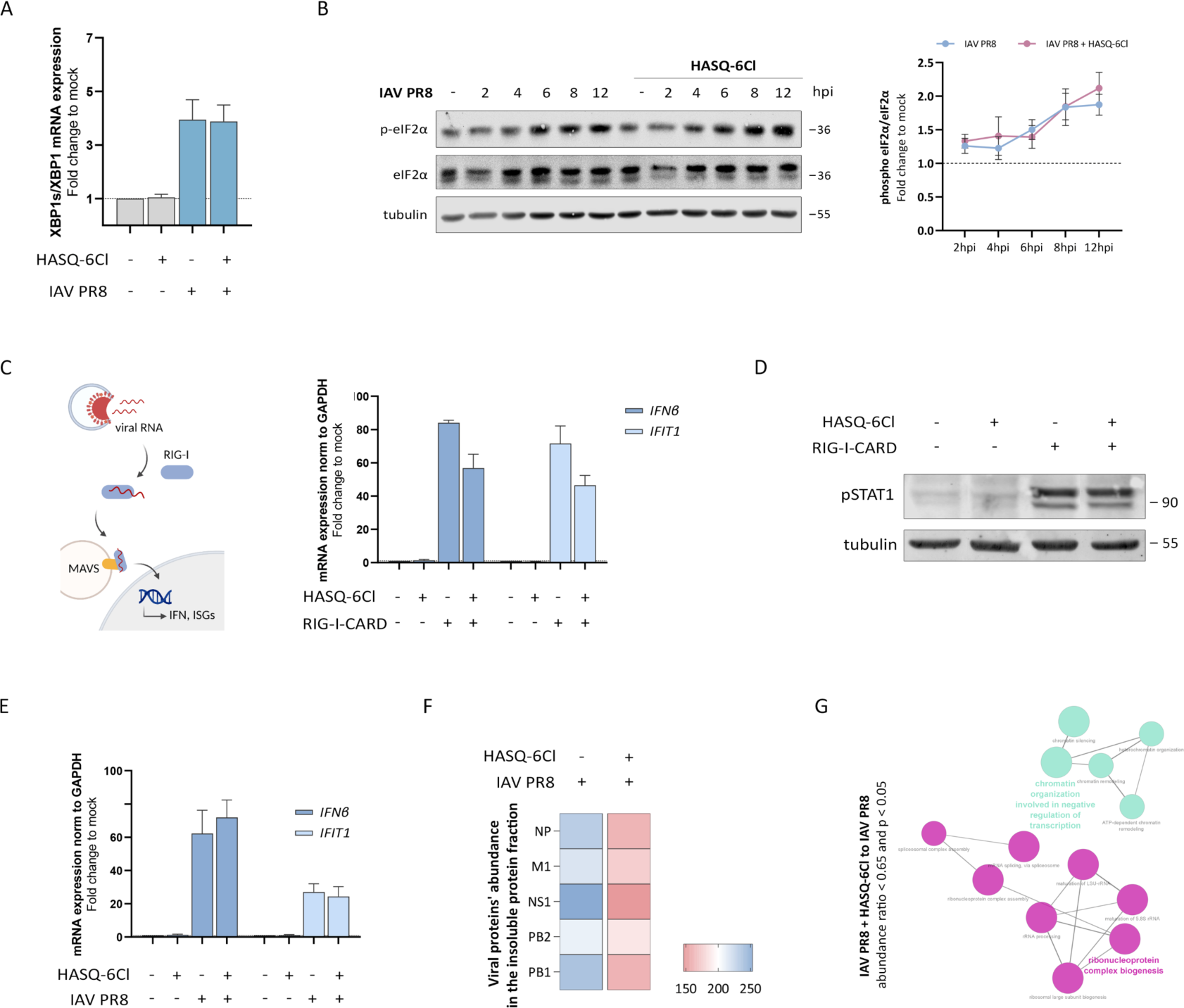
| Analysis of effect of HASQ-6Cl on the host cell’s immune response and UPR signaling. (A) RT-qPCR analysis of the splicing of XBP1 mRNA in A549 cells infected with IAV PR8 for 8 h in the absence or in the presence of HASQ-6Cl, in relation to mock-infected cells. Data represents the means ± SEM of three independent experiments. *p<0.05 in two-way ANOVA, with Bonferroni’s comparison test. (B) Western blot analysis of eIF2α phosphorylation in A549 cells infected with IAV PR8 at different times post infection in the absence or in the presence of HASQ-6Cl. Tubulin was used as internal control. (C-E) Analysis of the effect of HASQ-6Cl on the innate immune response. (C) RT-qPCR analysis of the IFNβ mRNA expression in A549 cells after stimulation with RIG-I-CARD for 10 h in the absence or in the presence of HASQ-6Cl, normalized to control. (D) Western blot analysis of pSTAT1 activation following A549 cells stimulation with RIG-I-CARD for 10 h in the absence or in the presence of HASQ-6Cl. (E) RT-qPCR analysis of the IFNβ mRNA expression in A549 cells after infection with IAV PR8 for 8 h in the absence or in the presence of HASQ-6Cl. Data represents the means ± SEM of three independent experiments. (F) Viral proteins’ abundance in the insoluble fraction of A549 cells infected with IAV PR8 in the presence or absence of HASQ-6Cl. Color gradient represents the relative abundance of each viral protein in comparison to other viral proteins’ abundances, considering both samples, from the lowest (red) to the highest (blue) value. (G) Characterization of the insoluble protein fraction of IAV PR8-infected cells in the presence of HASQ-6Cl. GO term analysis (using ClueGO) of the diminished insoluble proteins in cells infected with IAV PR8 after HASQ-6Cl treatment versus IAV PR8-infected cells (abundance ratio < 0.65 and p-value < 0.05 considering three biological replicates).

Another possibility might be that this compound could, by itself, induce the improvement of the cellular antiviral response against IAV. As most RNA viruses, IAV is mainly sensed by the RIG-I/MAVS antiviral signaling pathway [39], which culminates in the production of interferons (IFNs) or IFN-stimulated genes (ISG) that restrict the virus life cycle and warn the neighboring cells for the presence of the pathogen [40]. To determine whether HASQ-6Cl intensifies this antiviral signaling response, we analyzed the levels of IFNβ and the ISG IFN-induced protein with tetratricopeptide repeats 1 (IFIT1) in A549 cells upon transfection of RIG-I-CARD (a constitutively active form of RIG-I stimulates the RIG-I/MAVS pathway [41–43], in the absence or presence of HASQ-6Cl. Our results demonstrate that there is no significant difference between *IFNβ* or *IFIT1* production in the presence of HASQ-6Cl (**Fig. 5C**). In order to solidify these results, we have used the same experimental setup to analyze the activation of the signal transducer and activator of transcription 1 (STAT1, which signals IFNs produced by the neighboring cells) by western blotting using a specific antibody against the phosphorylated activated form of STAT1 (pSTAT1). As shown in **Fig. 5D**, there are no significant differences in the amount of pSTAT1 in the presence or absence of HASQ-6Cl upon stimulation with RIG-I-CARD. Finally, to prove that HASQ-6Cl has no influence on the host immune response against IAV, we did the same analysis in the context of IAV PR8 infection, in the presence and absence of the compound. Also in these conditions, there was no significant change in the levels of *IFNβ* and *IFIT1* mRNA (**Fig. 5E**). These results demonstrate that there is no specific interference of HASQ-6Cl with the host immune response dependent on MAVS signaling.

Altogether, up to this point, our results indicated that HASQ-6Cl’s action on physically preventing the formation of the virus-induced protein aggregates is the real cause of the delay in virus particle formation and the consequent decrease in virus titers. These results, hence, also solidify the hypothesis that the formation of these protein aggregates is in fact stimulated by the virus, as an important part of its infection cycle.

We further characterized by mass spectrometry the insoluble fraction of A549 cells upon 8 hpi with IAV PR8, after pre-treatment with HASQ-6Cl. We first performed comparative analyses of the abundances of each viral protein in the absence or in the presence of HASQ-6Cl during infection, considering three independent replicates. This analysis demonstrates that the proteins that were previously identified as present in the insoluble fraction of IAV PR8-infected cells (NP, NS1, PB1, PB2, M1) are less abundantly present when cells are treated with HASQ-6Cl prior to infection (**Fig. 5F** and Supplementary Table 5). Afterwards, to understand which host proteins were deregulated upon infection in the presence of HASQ-6Cl, we considered the proteins whose abundance ratio between IAV PR8-infected cells pre-treated with HASQ-6Cl (IAV PR8 + HASQ-6Cl) and IAV PR8-infected cells was below 0.65 with adjusted p-value < 0.05. A gene ontology analysis using both Cytoscape and STRING databases showed that most of the identified proteins are involved in the negative regulation of transcription and ribonucleoprotein complex biogenesis (**Fig. 5G** and Supplementary Table 6 and 7).

These results support our previous observations that indicate that the accumulation of both viral proteins and host proteins related to the ribonucleoprotein complex biogenesis (including mRNA splicing, ribonucleoprotein complex assembly or RNA processing) in insoluble aggregates plays an important role during IAV life cycle.

To further complete this study and confirm that the effect of HASQ-6Cl on infection is due to its protein disaggregation properties, we performed similar analyses upon infection with the non-segmented negative-strand RNA virus vesicular stomatitis virus (VSV). As shown in **Fig. 6A**, and in contrast to IAV, VSV does not induce a significant increase in the amount of insoluble proteins during infection. The quantification of the formation of VSV infectious particles by plaque assay in the presence of HASQ-6Cl rendered no significant differences when compared to the particles produced in the absence of this compound (**Fig. 6B**). These results hence suggest that the lack of effect of HASQ-6Cl on VSV infection is related to the absence of viral-induced protein aggregation and solidify our previous results on IAV, demonstrating that the observed accumulation of insoluble proteins is a specific characteristic of IAV infection which is essential for an efficient viral propagation.

**Figure 6.**
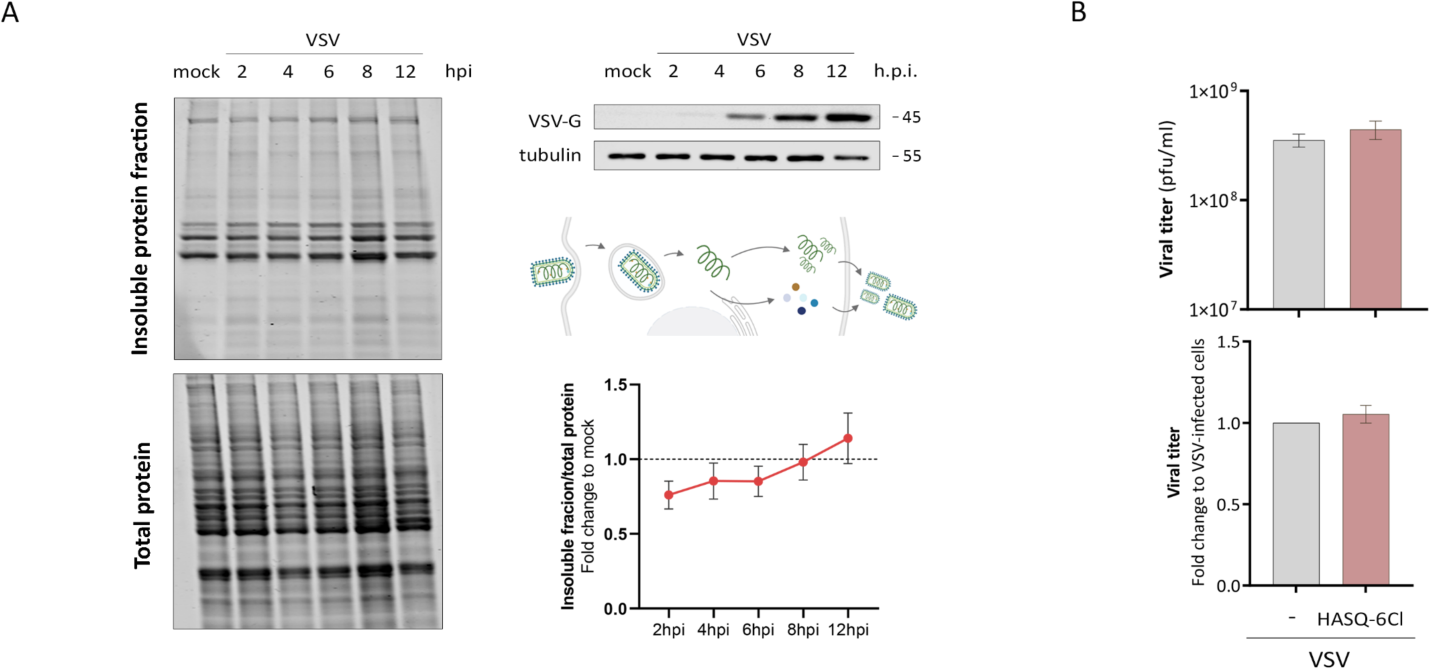
| Analysis of the accumulation of misfolded proteins in VSV-infected cells. (A) Characterization of the insoluble protein fraction of A549 cells infected with VSV at different times post infection. (B) VSV titers after treatment of A549 cells with HASQ-6Cl. Values were normalized to IAV PR8. Data represents means ± SEM of at least independent experiments, *p<0.05, ****p<0.0001 using student t-test.

### The disruption of virus-induced aggregates inhibits the proper assembly of new influenza virions

We have further analyzed the effect of the inhibition of the formation of protein aggregates on the expression of different IAV proteins throughout the whole infectious cycle (**Fig. 7A**). To that end, A549 cells were infected with IAV PR8 in the absence or presence of HASQ-6Cl, and the expression of IAV proteins was assessed at several times post infection by western blot.

**Figure 7.**
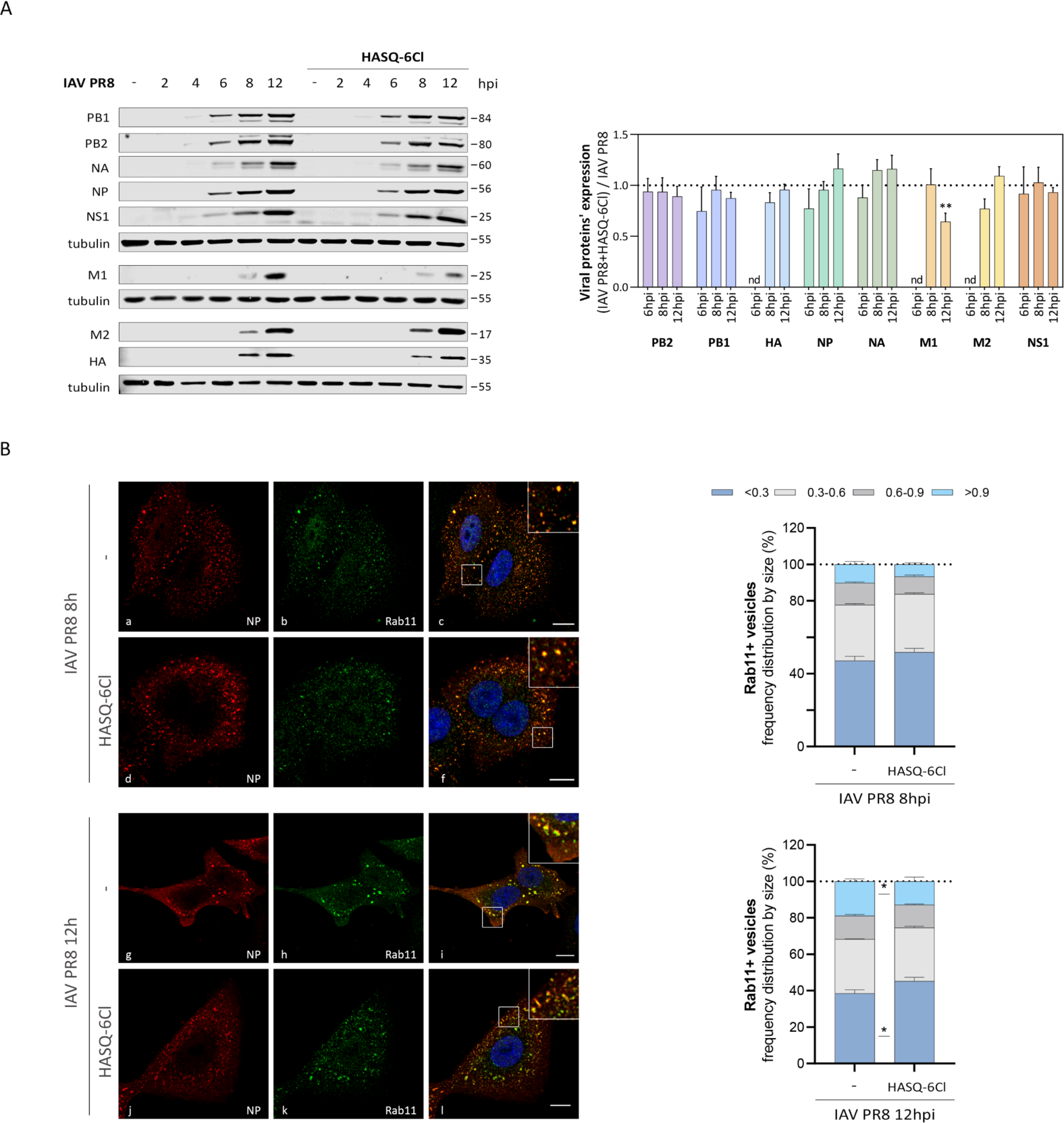
| HASQ-6Cl interferes with viral inclusions and impedes the proper assembly of new virus particles. (A) Western blot analysis of each of the viral proteins’ expression levels in IAV PR8-infected A549 cells at different times post infection, in the presence or absence of HASQ-6Cl. Tubulin was used as internal control. Quantification values were obtained after normalization to tubulin followed by the ratio between the intensity of a viral protein in a infected sample pre-treated with HASQ-6Cl (IAV PR8+HASQ-6Cl) and in an infected sample (IAV PR8), referred as (IAV PR8+HASQ-6Cl):IAV PR8. Data represents the means ± SEM of at least three independent experiments. **p<0.001 using student’s t-test. (B) Characterization of the viral inclusions’ size formed upon infection with IAV PR8 for 8 h or 12h in A549 cells, in the absence or in the presence of HASQ-6Cl. Confocal images of (a, d, g, j) viral NP, (b, e, h, k) Rab11 and (c, f, i, l) merge images. Bars represent 10 µm. Data represents the frequency distribution of viral inclusions by size as means ± SEM of three independent experiments, *p<0.05 using two-way ANOVA followed by Bonferroni’s multiple comparisons test.

As expected, most of the viral proteins, namely PB1, PB2, NP, are already detected after 6 hpi, while others, such as M1 and M2, are only expressed at later infection stages (**Fig. 7A**). Except for M1 at 12 hpi, none of the viral proteins’ expression was affected by the presence of HASQ-6Cl, indicating that neither the compound nor the presence of protein aggregates influences the general viral protein translation. Nevertheless, the formation of insoluble protein aggregates upon infection seems to be important for the correct expression, or at least stabilization, of M1. As M1 mediates viral assembly and budding at the plasma membrane of infected cells, by mediating the encapsidation of vRNPs into the membrane envelope [44–46], we hypothesize that the formation of these protein aggregates is somehow favoring the final assembly of the new virus particles.

The molecular details on how IAV eight-partite genome assembles are still not well understood. However recent data shows that vRNPs associate with a modified ER [47], to form insoluble liquid condensates, also referred as viral inclusions, at ER exit sites, through binding to the recycling endocysc marker Rab11a [27,47–49]. As infection progresses, viral inclusions augment in size. To understand whether the vesicular transport of vRNPs is being affected by the presence of HASQ-6Cl, A549 cells infected with IAV PR8, in the presence and absence of HASQ-6Cl, were stained against viral NP and host Rab11 and analyzed by immunofluorescence (**Fig. 7B**). Rab11-positive viral inclusion areas were visualized by confocal microscopy and quantified using the ImageJ software.

The vesicles were classified into groups according to size, based on previous reports [27,49,50]: we have set four intervals comprising vesicles with a size (1) up to 0.30 µm^2^, (2) 0.30-0.60 µm^2^, (3) 0.60-0.90 µm^2^ and (4) larger than 0.90 µm^2^. The frequency distribution of viral inclusions at 8 hpi is not significantly different in the presence or in the absence of HASQ-6Cl, although at 12 hpi there is an increase in the smaller and a decrease in the larger viral inclusions, indicating HASQ-6Cl is to some extent interfering with the formation of viral inclusions and, therefore, with the final assembly of new IAV virions.

## Discussion

During IAV particle formation, a large amount of viral proteins is synthesized in a relatively short time, whereby protein folding can become a limiting step for their active conformation and trafficking. Proteostasis disruption, including the one triggered by viral infections, may lead to the accumulation of misfolded proteins and the induction of ER stress, activating the UPR [51].

In this study we performed a detailed analysis of the activation and importance of each of the three (PERK-, ATF6- and IRE1-dependent) UPR pathways in the context of a single cycle of IAV infection. This approach differs from previous studies (which, as explained above, present somewhat contradicting data), that typically investigated the relevance of the UPR in IAV infection at fewer and mostly late time points post infection (24 or 48 h post infection) [16,20], often comprising cells in diverse stages/cycles of infection. We determined that, although the levels of eIF2α phosphorylation, alongside with the expression of ATF4, GADD34 and CHOP, increase upon infection, PERK is not essential for viral propagation, as the specific PERK inhibitor GSK-2656157 had no effect on the formation of new infectious virus particles. Furthermore, IAV infection in the presence of GSK-2656157 led to a similar pattern of eIF2α phosphorylation (and consequent ATF4 and GADD34 expression) to the one observed in non-treated infected cells. eIF2α phosphorylation during IAV infection may hence result from the activation of another signaling pathway independent of PERK. The integrated stress response (ISR) combines converging stress response pathways against diverse stimuli that culminate with eIF2α phosphorylation and its downstream signaling [52,53]. Besides PERK, other kinases are implicated in the ISR, namely protein kinase R (PKR), general control nonderepressible 2 (GCN2), and heme regulated inhibitor (HRI). As PKR has been shown to be activated in response to IAV infection [54], the observed changes are most likely due to the activation of this specific kinase. However, some IAV proteins can counteract this host defense factor [55–57], perhaps explaining, together with the ATF4/GADD34-driven negative feedback mechanism, why we observe a transient activation of this pathway.

On the other hand, inhibition of either ATF6 or IRE1 significantly affected the formation of new virus particles, suggesting that the activation of both pathways is important for infection. Although, due to technical constrains, we were not able to identify the specific activation of ATF6 during a single infection cycle, our results go along with others [16], where ATF6 cleavage was observed after 24 and 48 h of infection, implying a requirement of this pathway for a sustained infection after multiple cycles. On the other hand, our results opposed those that indicated that ATF6 did not play an important role upon IAV infection [20,21]. Nevertheless, one has to keep in mind that the used ATF6 inhibitor AEBSF, although widely applied as an ATF6 inhibitor, is a broad-spectrum serine protease inhibitor [58], which may mislead our conclusions. Further studies are then required to conclude on the importance of the UPR ATF6 pathway for IAV infection.

The IRE1 pathway activation was observed upon infection by the increase of IRE1 phosphorylation and the splicing of the downstream factor XBP1. Inhibition of this pathway using the specific inhibitor 4µ8C results in a significant decrease in the formation of new virus particles. Our results are in agreement and complementing to those obtained by Hassan *et al* [21], where the activation of the IRE1 pathway by XBP1 splicing was observed in HTBE cells, and by Schmoldt *et al* [20], that reported that a functional IRE1 is necessary for viral NP expression. The importance of the UPR IRE1/XBP1 pathway for IAV infection is most likely due to an XBP1-dependent upregulation of genes involved in phospholipids synthesis, chaperone expression and the activation of the ERAD machinery, that ultimately may lead to increased ER size and capacity to help viral protein folding and maturation. These results and the ones obtained with the ATF6 pathway may be correlated, as previous studies have shown that ATF6 can induce XBP1 mRNA [59,60], and that ATF6 seems to be required for full activation of XBP1 targets. Taken together, our results demonstrate the significance of both IRE1 and ATF6 pathways for IAV infection.

The stress-induced disruption of cellular proteostasis often results in the accumulation of insoluble proteins and toxic protein aggregates, which is proposed as a transversal hallmark of several pathological conditions [61,62]. In the context of a viral infection, the accumulation of proteins into aggregates can arise from the formation of specialized sites of viral replication and assembly (generally termed viral factories) or even as part of the host cell antiviral immune response [11,63,64]. In this study we investigated whether IAV induces the accumulation of insoluble protein aggregates and disturbs the host proteostasis. Our results show that, concomitantly to a high level of viral protein translation, IAV induces an increase in the amount of the cellular insoluble protein fraction and the formation of protein aggregates, mainly at the perinuclear area. This occurs at a point of infection that coincides with the activation of the UPR IRE1 pathway and with a high viral protein translation rate, while host protein synthesis is considerably decreased [36].

To elucidate the composition, origin and significance of the observed IAV-induced protein aggregation, we analyzed the insoluble protein fractions by LC-MS/MS. IAV proteins NP, NS1, PB1, PB2 and M1 accumulate in the insoluble protein fraction, as well as host proteins associated to mechanisms of protein complex assembly and localization, mRNA processing and protein translation. A recent study has demonstrated that, during IAV infection, several host proteins undergo changes in solubility, and found that several viral proteins, essentially vRNP components, become strongly insoluble with infection [65], strengthening our results.

Additionally, a previous study found several proteins belonging to the above-mentioned pathways immunoprecipitated with IAV H7N9 NP [66]. It is possible that NP recruits these proteins to enhance viral transcript translation and that they generate large insoluble complexes. Our results may also indicate that these translation related processes are deregulated upon the export of viral ribonucleoproteins from the nucleus and the viral host-shutoff to impede host proteins to be fully synthesized, which ultimately may lead to ribosome stalling and accumulation of host aberrant insoluble proteins.

To better understand the biological relevance of the accumulation of insoluble protein aggregates in the context of IAV infection, we used a tool to inhibit this aggregation, the hybrid chemical compound HASQ-6Cl (named 6c in [38]). The combination of quinolines and steroids in one single chemical entity, generates this new hybrid molecule, capable of interacting with protein aggregates through π-π (quinoline fragment), hydrophobic (steroid fragment) and hydrogen bonding interaction. Both fragments of HASQ-6Cl have important features to be able to interact with β-sheets inhibiting its consequent aggregation process. Such versatility is critical to interact with protein aggregates of yet unknown origin, such as those observed in this work. The design strategy of HASQ-6Cl followed a “framework combination” approach, avoiding the use of cleavable linkers, to retain the molecule integrity within the cellular media. Our results demonstrate that HASQ-6Cl decreases the IAV-induced protein aggregation level. Importantly, we have shown that HASQ-6Cl interferes with the virus life cycle at the viral protein production phase, which coincides with the protein aggregates formation, and consequently induces about 50% decrease in the production of infectious viral particles. This effect is likely due to the diminished formation of the virus-induced protein aggregates, as no interference of HASQ-6Cl on the splicing of XBP1, and hence on IRE1 activation, nor on the RIG-I/MAVS antiviral signaling were observed.

Mass spectrometry analysis of the insoluble fractions isolated upon IAV infection in the presence of HASQ-6Cl revealed that the viral proteins that were previously identified as present in the insoluble fraction of infected cells are less abundant in these conditions. Indeed, the presence of HASQ-6Cl during infection decreases the abundance of several viral proteins with different roles during the infection cycle, namely vRNP components (PB1, PB2 and NP), NS1 and M1, that were shown here, and by others [65] to become more insoluble upon infection with IAV PR8.

There were also changes observed at the level of the host proteins, with most of the identified proteins being involved in the negative regulation of transcription and ribonucleoprotein complex biogenesis. These results support the hypothesis that the accumulation of both viral proteins and host proteins in insoluble aggregates, or changes in its solubility, plays an important role during IAV infection. Furthermore, it suggests that HASQ-6Cl might have a multiple and broad targeting within cells to prevent the accumulation of proteins in the insoluble fraction and, therefore, restricting infection.

The proper assembly and budding of new virions require the intra-cellular transport of progeny vRNPs from the nucleus to the plasma membrane. This process is dependent on the formation of liquid viral inclusions that are hubs for the eight vRNPs close to ER exit sites, and their formation and/or transport to the plasma membrane depends on the ER-derived vesicles containing Rab11 [67–70]. Recently, it was shown that nucleozin, a well-studied vRNP pharmacological modulator, can affect vRNP solubility in a Rab11-dependent manner, acting by hardening IAV inclusions to prevent efficient replication [65]. Our results show that the formation of these viral inclusions can be impacted HASQ-6Cl by decreasing their size, which is ultimately reflected in a decrease in the number of infectious IAV particles produced after a single replication cycle. Targeting this mechanism may be of particular importance as, besides being crucial for the correct assembly of every genome segment to form a fully infectious viral particle, it is also decisive for the genetic reassortment upon a co-infection of different IAV strains from distinct hosts and the emergence of novel viruses with pandemic potential [71].

To complete our studies, and determine whether HASQ-6Cl, and its consequent inhibition of protein aggregation, affected the production of specific viral proteins, we assessed their expression at several times post infection in the absence or presence of the chemical compound. We detected no difference in the production of viral proteins in the presence of HASQ-6Cl, with the exception of M1. M1 is the most abundant protein in virions and mediates viral assembly and the budding of vRNPs at the plasma membrane of infected cells [44,45]. This protein has different roles at different stages of the infection, and it is speculated that it can change its conformational and oligomerization state depending on its functional state [72]. One hypothetical function of the M1 oligomer is the shielding of newly synthesized vRNA/vRNPs during transport through the host cell’s cytosol after nuclear export. One can then further hypothesize that the presence of HASQ-6Cl can in some extent prevent the oligomerization of M1, indirectly manipulating its function during assembly through vRNPs destabilization.

The formation of these protein aggregates and their relevance for viral propagation seems to be specific for IAV, or at least not common to all RNA viruses, as we have shown that VSV infection does not lead to an accumulation of insoluble proteins, and that the presence of HASQ-6Cl does not affect the production of infectious VSV particles.

Overall, our findings demonstrate that IAV manipulates the host cellular processes by activating the UPR and by inducing the accumulation of insoluble viral proteins and host proteins that are generally related to RNA processing. The formation of these aggregates is beneficial for the virus and seems to be required for the correct assembly of viral particles. Interfering with UPR pathways or chemically avoiding the assembly of such aggregates is sufficient to hinder viral propagation. Our results have, hence, uncovered specific IAV-host interaction mechanisms that should be further explored for the development of novel host-directed IAV antiviral therapies.

## Methods

### Cells, plasmids and transfection

A549, MDCK, HEK293T, Vero and HeLa expressing a HSP27:GFP reporter (HeLa HSP27:GFP) cells were cultured in Dulbecco’s modified Eagle’s medium (DMEM) high glucose (4,5 g/L) supplemented with 10% fetal bovine serum (FBS) and 100U/mL of penicillin and 100 mg/mL streptomycin (all from GIBCO) and incubated at 37°C in a humidified atmosphere containing 5% CO_2_. All cells were regularly tested for mycoplasma contamination. When indicated, A549 cells were transfected with GFP-RIG-I-CARD by 10h incubation with Lipofectamine 3000 (Invitrogen) according to manufacturers’ protocol.

### Virus stock preparation

Reverse-genetics derived influenza A/Puerto Rico/8/34 (IAV PR8; H1N1) were rescued in 293T cells using a pDUAL plasmid system. This virus was produced in HEK293T cells upon transfection of eight plasmids encoding the gene segments of IAV PR8 using PEI transfection reagent in DMEM high glucose (4.5 g/L) with pyruvate (0.11 g/L), supplemented with glutamine, without FBS or antibiotics. 24 h post transfection, the medium was changed to 2 mL of serum-free media supplemented with 1% P/S, with 0.14% BSA and 1 µg/mL trypsin-TPCK. After two days of incubation under the same growing conditions, the supernatant was collected and centrifuged at 3000 rpm for 5 min. Clarified supernatant with low number of viruses (P0) was stored at −80°C and further quantified by plaque assay. To prepare virus stocks, MDCK cells were infected with P0 aliquots in serum-free media (SFM) supplemented with 0.14% BSA and 1 µg/mL trypsin-TPCK, with a MOI of 0.01. After virus adsorption for one hour, cells were cultured for two days at 37 °C, 5% CO_2_. After centrifugation for 5 min at 3000 rpm, clarified supernatants were aliquoted and stored at −80 °C. Virus stock or samples titers were determined by plaque assay.

Vesicular stomatitis virus (VSV) Indiana strain parental stock was kindly provided by Dr. Jonathan Kagan (Boston Children’s Hospital, Harvard Medical School). All virus stocks were propagated in highly confluent Vero cells, using a MOI of 0.001. The virus inoculum was diluted in SFM and after virus adsorption for one hour, an incubation of 18 h ensued. A centrifugation for 5 min at 500 g, 4 °C was carried out to collect the clarified supernatant. Then, an ultracentrifugation of the supernatant for 90 min at 24000 rpm, 4 °C followed and, once the supernatant was discarded, 0.35 mL of NTE buffer (0.1M NaCl, 1mM EDTA, 0.1M Tris pH 7.4) was added and incubated on ice, overnight. The next day, 4 mL of sucrose 10% NTE (ice-cold) was overlayed with 1 mL of virus suspension and ultracentrifuged for 60 min at 40500 rpm, 4 °C. The supernatant was removed, and the pellet was incubated with 0.5 mL of NTE buffer on ice, overnight. Finally, the virus suspension was aliquoted and stored at −80 °C. Virus stock or samples titers were determined by plaque assay.

### Plaque assay and infection experiments

To quantify IAV, MDCK cells were cultured with 10-fold dilutions of virus suspension and allowed to absorb for 1h. Cells were then cultured in 50% avicel-containing SFM supplemented with 0.14% BSA and 1 µg/mL trypsin-TPCK for 1.5 to 2 days. Cellular monolayers were fixed in 4% paraformaldehyde and stained with 0.1% toluidine blue. To quantify VSV, Vero cells were used and the avicel-containing SFM media were in this case supplemented with 1% FBS and the protocol ran for 1 day.

To perform infection, A549 cells were seeded to achieve a confluency of 80% at the time of the infection. The day after, cells were washed with PBS and infected with IAV PR8 or VSV at a MOI of 3, prepared in SFM. After one hour, cells were overlaid with DMEM supplemented with 20% FBS and 1% P/S and incubated for the desired times at 37 °C in 5% CO_2_. In the case of samples used to determine viral production, cells were infected and after 1 h incubation, the supernatant was removed, cells were rinsed with acid wash buffer 1x (135 mM NaCl, 10 mM KCl, 40 mM citric acid, pH 3), washed in PBS and incubated in SFM supplemented with 0.14% BSA, and 1 µg/mL trypsin-TPCK (for IAV only), for up to 16 hpi.

### Unfolded protein response inhibitors

PERK inhibitor GSK-2656157 (BIOV9466-5, VWR), IRE1 inhibitor 4µ8C (412512, Calbiochem) or ATF6 inhibitor AEBSF (#5175, Tocris Bioscience) were added to the cells, at the indicated concentrations, 1 h prior to infection. Different concentrations of each inhibitor were tested to check for cell viability to define which one to use. The number of the infectious particles formed were normalized to the cell viability in each condition.

### Experiments with the HASQ-6Cl compound

The HASQ-6Cl compound was solubilized in 100% ethanol (EtOH) to obtain a stock concentration of 100 mM. The cytotoxicity of HASQ-6Cl and EtOH was assessed by MTT cell viability assay. To do that, cells were seeded into a 96-well plate at a density of 8×10^3^ cells/plate and allowed to adhere for 24 h at growing conditions. The day after, cells were treated with various concentrations of the compound (100, 50, 25 and 10 µM) or the correspondent EtOH percentage (1, 0.5, 0.25 and 0.1%) and incubated for 30 h at 37 °C in a CO_2_ incubator. Compound-containing solutions were sterilized using a 0.2 µm filter. Cells were then washed twice with PBS and incubated to DMEM with 10% MTT (working solution 5 mg/mL in phosphate buffer) for 2 h at growing conditions. Lastly, this medium was removed, and the formazan crystals formed were solubilized using DMSO for 10 min at room temperature and the intensity was quantified at 575 nm. Untreated cells were used as control and the blank value was subtracted to all conditions. Further experiments were performed using 50 µM of HASQ-6Cl. The compound was added to the cells 12h before infection.

### Immunocytochemistry and microscopy analyses

Cells grown in 12ømm glass coverslips were fixated for 20 min with 4% paraformaldehyde in PBS, pH 7.4, followed by permeabilization with 0.2% triton X-100 for 10 min, blocking with 1% BSA for 10 min, and immunostaining with the indicated primary and secondary antibody for 1 h each. This procedure was done at room temperature, with cells being washed three times between each step. Primary antibodies used include anti-nucleoprotein (ab20343, Abcam), and anti-Rab11 (sc-166912, Santa Cruz Biotechnology). The fluorophores used include Alexa488 (A21206 and A21202, Invitrogen) or TRITC (711-025-152, Jackson ImmunoResearch). When needed, staining with Proteostat Aggresome Dectetion kit (Enzo Life Sciences International) was performed for 30 min. Cells were additionally incubated with Hoechst dye for 2 min before being mounted in slides, using Mowiol 4-88 (AppliChem Inc.) containing propyl gallate (Sigma-Aldrich). Confocal images were acquired using a Zeiss LSM 880 confocal microscope (Carl Zeiss) and a Plan-Apochromat 63× and 100×/1.4 NA oil objectives, a 561 nm DPSS laser and the argon laser line 488 nm (BP 505–550 and 595–750 nm filters). Images were processed using ZEN Black and ZEN Blue software (Carl Zeiss).

Quantification of the NP intensity in PR8-infected A549 cells was performed after sketching the region of interest (ROI) of the nucleus of each cell and obtaining the correspondent intensity mean gray value using the Zeiss Blue Software for image processing (Carl Zeiss). Characterization of viral inclusions, positive for Rab11, by size in infected cells was performed using Fiji/ImageJ software (NIH). The vesicles were divided into groups according to size, based on previous reports [27,48,50]. In uninfected cells, the areas for viral inclusions are consistent with values <0.15 µm^2^ [50]. With infection, the frequency distribution from inclusions measuring 0.15-0.30 µm^2^ and bigger than 0.3 µm^2^ augmented significantly in relation to non-infected cells [49]. As we propose to compare the areas in infected cells only, we have set four intervals comprising viral inclusions with a size (1) up to 0.30 µm^2^, (2) 0.30-0.60 µm^2^, (3) 0.60-0.90 µm^2^ and (4) larger than 0.90 µm^2^.

### Isolation of the insoluble protein fraction

To obtain total protein extracts, cell pellets were resuspended in Empigen Lysis Buffer (ELB) (0.5% Triton X-100, 50 mM HEPES, 250 mM NaCl, 1 mM DTT, 1 mM NaF, 2 mM EDTA, 1 mM EGTA, 1 mM PMSF, 1 mM Na_3_VO_4_ supplemented with a cocktail of protease inhibitors). Protein extracts were then sonicated, and centrifuged 20 min at 200 g at 4 °C. In the end, supernatants were kept for the next phase of total protein quantification. During all procedures, cells remained on ice to avoid the activity of proteases. Quantification of total protein was performed using Pierce™ Bovine Serum Albumin Protein Assay Kit (Thermo Scientific), following the manufactureŕs instructions.

To isolate the insoluble protein fraction, 100 μg of total protein was diluted in ELB to reach a final volume of 100 µL. Samples were centrifuged for 16,000 g, 20 min at 4 °C to obtain the insoluble fraction. The resulting pellet was solubilized in ELB supplemented with 20% NP-40 (10%). Samples were sonicated for three cycles of five seconds (0.5 cycles with amplitude at 50-60%). After sonication cycles, samples underwent another centrifugation round (16,000 g, 20 min at 4 °C). The supernatant was removed, the pellet was resuspended in ELB, and after adding the loading buffer, samples were denatured at 95 °C for 5 min and run into an SDS-PAGE. In parallel, 10 μg of total protein extracts were run as loading controls. Finally, the gel was stained with BlueSafe solution for at least 30 min and, after destained with distilled water, gels were scanned using Odyssey Infrared Imaging System (LI-COR Biosciences). The relative amount of insoluble protein for each sample was calculated by dividing the intensities of the insoluble protein extracts by the intensities of the corresponding total protein extracts.

### Gel Electrophoresis and Immunoblotting

Cells were pelleted and washed before being resuspended in ELB supplemented with protease-inhibitors mix. Samples were then incubated on a rotary mixer for 15 min at 4 °C and, after sonication using Clifton SW3H 38kHz bath (5 cycles 30 s on; 30 s off), the incubation was repeated. After clearing by centrifugation (17000 g, 15 min at 4 °C, the supernatant was collected, and protein concentration was determined using Pierce™ Bovine Serum Albumin (BCA) Protein Assay Kit (Thermo Scientific) . Protein samples were separated by SDS-PAGE on 10 or 12.5% polyacrylamide gels, alongside with a pre-stained protein marker (GRS Protein Marker Multicolour Tris-Glicine 4∼20%, Grisp), wet transferred to nitrocellulose (PROTAN®), and analyzed by immunoblotting.

Immunoblots were processed using specific primary antibodies and IRDye 800CW or IRDye 680RD secondary antibodies (#926-32211 and #926-68070, LI-COR Biosciences,). Primary antibodies used include anti-nucleoprotein (ab20343, Abcam, Cambridge, UK), anti-hemagglutinin (sc-52025, Santa Cruz Biotechnology), anti-neuraminidase (GTX636323, Genetex), anti-M1 (sc-69824, Santa Cruz Biotechnology), anti-M2 (sc-32238, Santa Cruz Biotechnology), anti-IRE1 (14C10 (3294), Cell Signaling Technology), anti-IRE1 (pS724) (ab48187, Abcam), anti-CHOP (L63F7, Cell Signaling Technology), anti-eIF2α (D-3) (sc-133132, Santa Cruz Biotechnology), anti-phospho-eIF2α (S1) (ab32157, Abcam), anti-ATF4 (BS6475, Bioworld), anti-GADD34 (PA5-30486, Invitrogen), anti-CHOP (L63F7, Cell Signaling Technology), anti-PERK (C33E10, #3192, Cell Signaling Technology), p-STAT1 (Y701, BD Biosciences) and anti-tubulin (T9026, Sigma-Aldrich). For quantification, immunoblots were scanned with Odyssey CLx (LI-COR Biosciences) and processed using the volume tools from Image Studio Lite Ver 5.2 software (LI-COR Biosciences).

### RNA extraction, cDNA synthesis and quantitative real-time polymerase chain reaction

Total RNA was isolated using NZYol reagent according to manufacturer’s instructions and quantified using DS-11 spectrophotometer (DeNovix Inc.), cDNA synthesis was obtained using 0.5 to 1 μg RNA, after a pre-treatment with DNAse (Thermo Scientific), using Revert Aid Reverse Transcriptase (Thermo Scientific) and Oligo-dT15 primer (Eurofins Genomics) following manufacturer’s protocol.

When needed, primer sequences were designed using Beacon Designer 7 (Premier Biosoft). The oligonucleotides used for human XBP1 were 5’-TTACGAGAGAAAACTCATGGCC-3’ and 5’-GGGTCCAAGTTGTCCAGAATGC-3’; for XBP1 spliced were 5’-CTGAGTCCGAATCAGGTGCAG-3’ and 5’-ATCCATGGGGAGATGTTCTGG-3’; for human IFN-β were 5’-TGGCACAACAGGTAGTAGGC-3’ and 5’-TGGAGAAGCACAACAGGAGAG-3’; and human GAPDH were 5’-AAGGTGAAGGTCGGAGTC-3’ and 5’-GGGTGGAATCATATTGGAACAT-3’ (Eurofins Genomics). GAPDH was used as a reference gene. RT-qPCR was performed in duplicate using 2x SYBR Green qPCR Master Mix (Low Rox) (Bimake), cDNA samples were diluted 1:10 and the concentration of each primer was 250 nM, with the exception for IFN-β in which we use undiluted samples and a primer concentration of 500 nM. Reactions were run on Applied Biosystems 7500 Real Time PCR System (Applied Biosystems). The thermocycling reaction was done by heating at 95 °C during 3 min, followed by 40 cycles of a 12 s denaturation step at 95 °C and a 30 s annealing/elongation step at 60 °C. The fluorescence was measured after the extension step using the Applied Biosystems software (Applied Biosystems). After the thermocycling reaction, the melting step was performed with slow heating, starting at 60 °C and with a rate of 1%, up to 95 °C, with continuous measurement of fluorescence. Data analysis was performed using the quantitative 2−ΔΔCT method.

### Statistical Analyses

Statistical analysis was performed with Graph Pad Prism 9 (GraphPad Software). Quantitative data were attained from at least three independent experiments and represent the means ± standard error mean (SEM). To determine the statistical significance among the experimental groups the two-way ANOVA followed by Dunnett’s multiple comparison tests were applied; comparison between two groups were made by Student’s t test. P values of ≤ 0.05 were considered as significant.

### LC-MS/MS analyses

Each sample was processed for proteomics analysis following the procedure described in [73] with implementation of the solid-phase-enhanced sample-preparation (SP3) detailed in [74]. Enzymatic digestion was performed with Trypsin/LysC (2 µg) overnight at 37°C at 1000 rpm. Protein identification and quantitation was performed by nanoLC-MS/MS. This equipment is composed by an Ultimate 3000 liquid chromatography system coupled to a Q-Exactive Hybrid Quadrupole-Orbitrap mass spectrometer (Thermo Scientific, Bremen, Germany). Samples were loaded onto a trapping cartridge (Acclaim PepMap C18 100Å, 5 mm x 300 µm i.d., 160454, Thermo Scientific) in a mobile phase of 2% ACN, 0.1% FA at 10 µL/min. After 3 min loading, the trap column was switched in-line to a 50 cm by 75μm inner diameter EASY-Spray column (ES803, PepMap RSLC, C18, 2 μm, Thermo Scientific, Bremen, Germany) at 300 nL/min. Separation was generated by mixing A: 0.1% FA, and B: 80% ACN, with the following gradient for total protein fraction: 5 min (2.5% B to 10% B), 120 min (10% B to 30% B), 20 min (30% B to 50% B), 5 min (50% B to 99% B) and 10 min (hold 99% B). Subsequently, the column was equilibrated with 2.5% B for 17 min and a specific gradient for the insoluble fractions: 5 min (2.5% B to 10% B), 30 min (10% B to 30% B), 50 min (30% B to 50% B), 45 min (50% B to 99% B) and 30 min (hold 99% B). Data acquisition was controlled by Xcalibur 4.0 and Tune 2.9 software (Thermo Scientific, Bremen, Germany).

The mass spectrometer was operated in data-dependent (dd) positive acquisition mode alternating between a full scan (m/z 380-1580) and subsequent HCD MS/MS of the 10 most intense peaks from full scan (normalized collision energy of 27%). ESI spray voltage was 1.9 kV. Global settings: use lock masses best (m/z 445.12003), lock mass injection Full MS, chrom. peak width (FWHM) 15 s. Full scan settings: 70 k resolution (m/z 200), AGC target 3e6, maximum injection time 120 ms. dd settings: minimum AGC target 8e3, intensity threshold 7.3e4, charge exclusion: unassigned, 1, 8, >8, peptide match preferred, exclude isotopes on, dynamic exclusion 45 s. MS2 settings: microscans 1, resolution 35k (m/z 200), AGC target 2e5, maximum injection time 110 ms, isolation window 2.0 m/z, isolation offset 0.0 m/z, spectrum data type profile.

The raw data was processed using Proteome Discoverer 2.4.0.305 software (Thermo Scientific) and searched against the UniProt database for the Homo sapiens Proteome 2020_01 (74,811 sequences) and the UniProt database for influenza A virus (strain A/Puerto Rico/8/1934 H1N1) 2020_04. The Sequest HT search engine was used to identify tryptic peptides. The ion mass tolerance was 10 ppm for precursor ions and 0.02 Da for fragmented ions. The maximum allowed missing cleavage sites was set to 2. Cysteine carbamidomethylation was defined as constant modification. Methionine oxidation and protein N-terminus acetylation were defined as variable modifications. Peptide confidence was set to high. The processing node Percolator was enabled with the following settings: maximum delta Cn 0.05; decoy database search target FDR 1%, validation based on q-value. Protein label free quantitation was performed with the Minora feature detector node at the processing step. Precursor ions quantification was performing at the processing step with the following parameters: unique plus razor peptides were considered for quantification, precursor abundance was based on intensity, normalization mode was based on total peptide amount, protein ratio calculation was pairwise ratio based, imputation was not performed, hypothesis test was based on t-test (background based).

The mass spectrometry proteomics data have been deposited to the ProteomeXchange Consortium [75] via the PRIDE [76] partner repository with the dataset identifier PXD043310. Gene ontology analysis of LC-MS/MS data was performed using Cytoscape (ClueGO app) or STRING online database.

## Acknowledgments

We would like to thank Dr Catarina Almeida for kindly providing anti-PERK and anti-GADD34 antibodies, the IRE1 inhibitor 4μ8C, the PERK inhibitor GSK-2656157 and XBP1 primers for RT-qPCR. We would also like to thank Dr Bruno Neves for kindly providing anti-CHOP, anti-IRE1, and anti-phosphorylated IRE1 antibodies, as well as the ATF6 inhibitor AEBSF. We also thank Dr Friedemann Weber for kindly providing the GFP-RIG-I-CARD plasmid. Dr. Sandra Vieira is also thanked for kindly providing the anti-Rab11 antibody. We also want to express our gratitude to Dr. Hugo Osório for his help in the mass spectrometry data acquisition and subsequent analysis. Some of the figures were created using BioRender.com.

This work was supported by the Portuguese Foundation for Science and Technology (FCT): POCI-01-0145-FEDER-031378, POCI-01-0145-FEDER-016630, POCI-01-0145-FEDER-029843, 2022.06064.PTDC, CEECIND/03747/2017, CEECIND/00284/2018, SFRH/BD/137851/2018, UIDB/04501/2020, and UIDP/50006/2020, under the scope of the Operational Program “Competitiveness and internationalization” (COMPETE 2020), in its FEDER/FNR component. It was also funded by the CCDRC, FEDER: pAGE - CENTRO-01-0145-FEDER-000003 and CENTRO-01-0246-FEDER-000018. The work was also funded by the European Union’s Horizon 2020 research and innovation program (grant H2020-WIDESPREAD-2020-5 ID-952373, for D.R. and A.R.S; and European Research Council (ERC) grant agreement No. 101001521, for M.J.A.).

## Declaration of interests

The authors declare no competing interests.

## Supplementary files

**Supplementary Figure 1.**
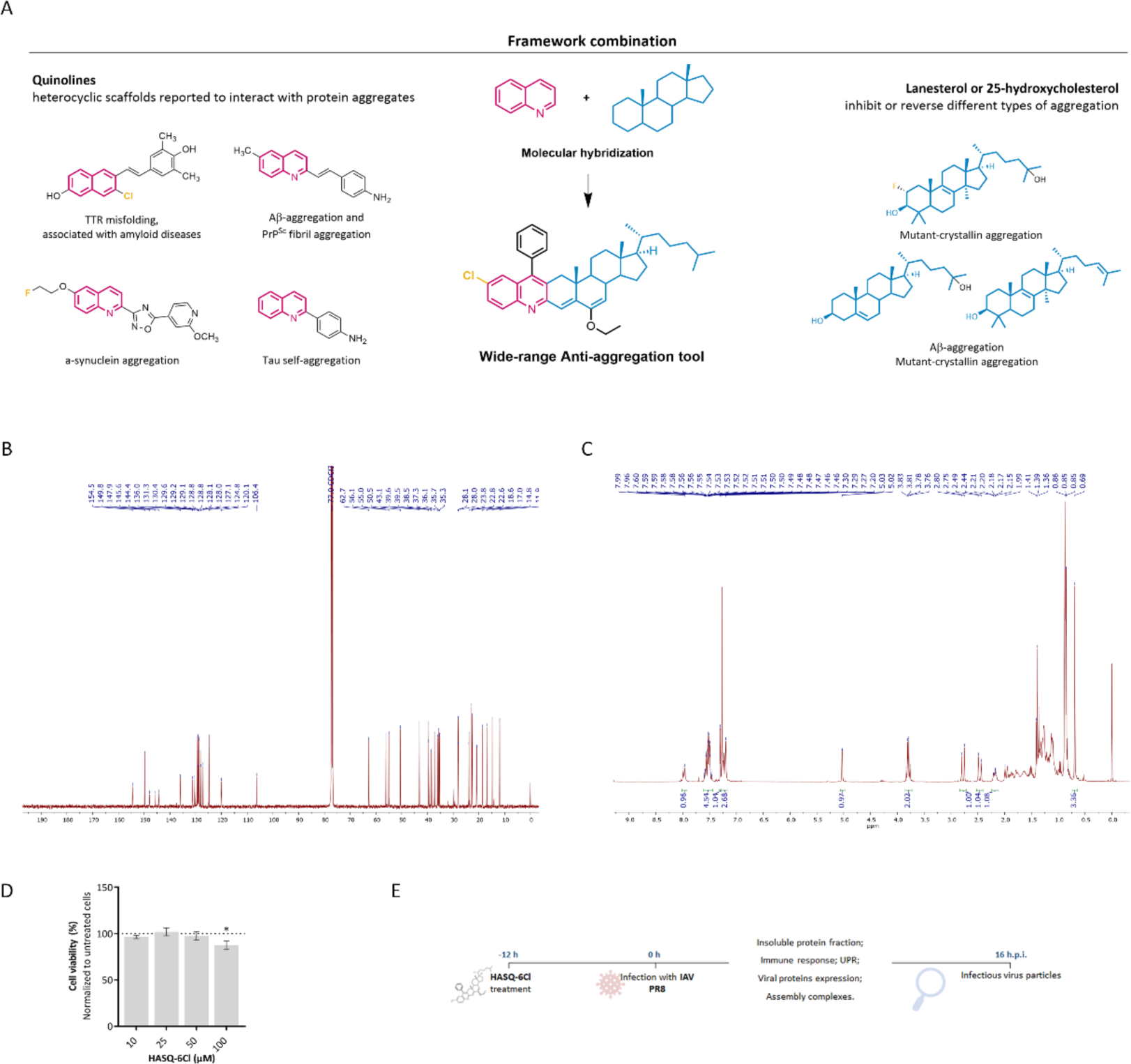
| HASQ-6Cl design strategy and toxicity. (A) Seminal examples of quinolines (in pink) and steroids (in blue) which were demonstrated to be capable to inhibit or reverse different types of protein aggregation. The combination of quinolines and steroids in one single new chemical entity, generates a new hybrid molecule, HASQ-6Cl, capable to interact with protein aggregates through π-π (quinoline fragment), hydrophobic (steroid fragment) and hydrogen bonding interactions. Schematic representation of the selected design strategy for HASQ-6Cl by framework combination. (B) ^13^C-NMR of compound HASQ-6Cl (CDCl_3_, 125 MHz). (C) ^1^H-NMR of compound HASQ-6Cl (CDCl_3_, 300 MHz). (D) Cell viability evaluated by MTT assay in A549 cells after 30 h of exposure to 10, 25, 50 and 100 µM HASQ-6Cl. Data shows the percent of viable cells normalized to untreated control (means ± SEM of 8 independent experiments). *p<0.05 using student’s *t-test*. (E) Schematic representation of experimental design used.

**Supplementary Table 1.** | Identification of influenza virus (strain A/Puerto Rico/8/1934 H1N1) proteins in the insoluble protein fractions of infected A549 cells at 8 hpi (n=3) Proteins were considered to be present in the insoluble protein fractions in infected samples when having the number of peptides and unique peptides ≥ 2, an abundance ratio (IAV PR8) / (mock) ≥ 1.5, and an abundance ratio adjusted *p-value* (IAV PR8) / (mock) ≤ 0.05.

| Accession | Description |  | # Peptides | # Unique Peptides | Abundance Ratio (IAV PR8):(mock) | Adj. p-Value (IAV PR8):(mock) | Found in Sample: mock #1 | Found in Sample: mock #2 | Found in Sample: mock #3 | Found in Sample: IAV PR8 #1 | Found in Sample: IAV PR8 #2 | Found in Sample: IAV PR8 #3 |
| --- | --- | --- | --- | --- | --- | --- | --- | --- | --- | --- | --- | --- |
| N0BQ24 | Polymerase basic protein 1 | PB1 | 2 | 2 | 100 | 0.000 | Not Found | Not Found | Not Found | Not Found | Not Found | High |
| A0A385P512 | Matrix protein 1 | M1 | 6 | 6 | 15.867 | 0.000 | Not Found | Peak Found | Peak Found | High | High | High |
| P03466 | Nucleoprotein | NP | 20 | 17 | 13.254 | 0.000 | Peak Found | Peak Found | Peak Found | Peak Found | High | High |
| I6TWM9 | Non-structural protein 1 | NS1 | 11 | 11 | 8.592 | 0.000 | Peak Found | High | Peak Found | Peak Found | High | High |
| A4ZNP4 | Polymerase basic protein 2 | PB2 | 12 | 12 | 6.403 | 0.001 | Peak Found | Peak Found | Peak Found | Peak Found | High | High |
| A4ZNP6 | Polymerase acidic protein | PA | 2 | 2 | 2.953 | 0.244 | Not Found | Peak Found | Not Found | Not Found | Not Found | High |
| T2AA52 | Hemagglutinin | HA | 3 | 3 | 1.216 | 0.892 | Not Found | Peak Found | Peak Found | Peak Found | High | High |

**Supplementary Table 2.**
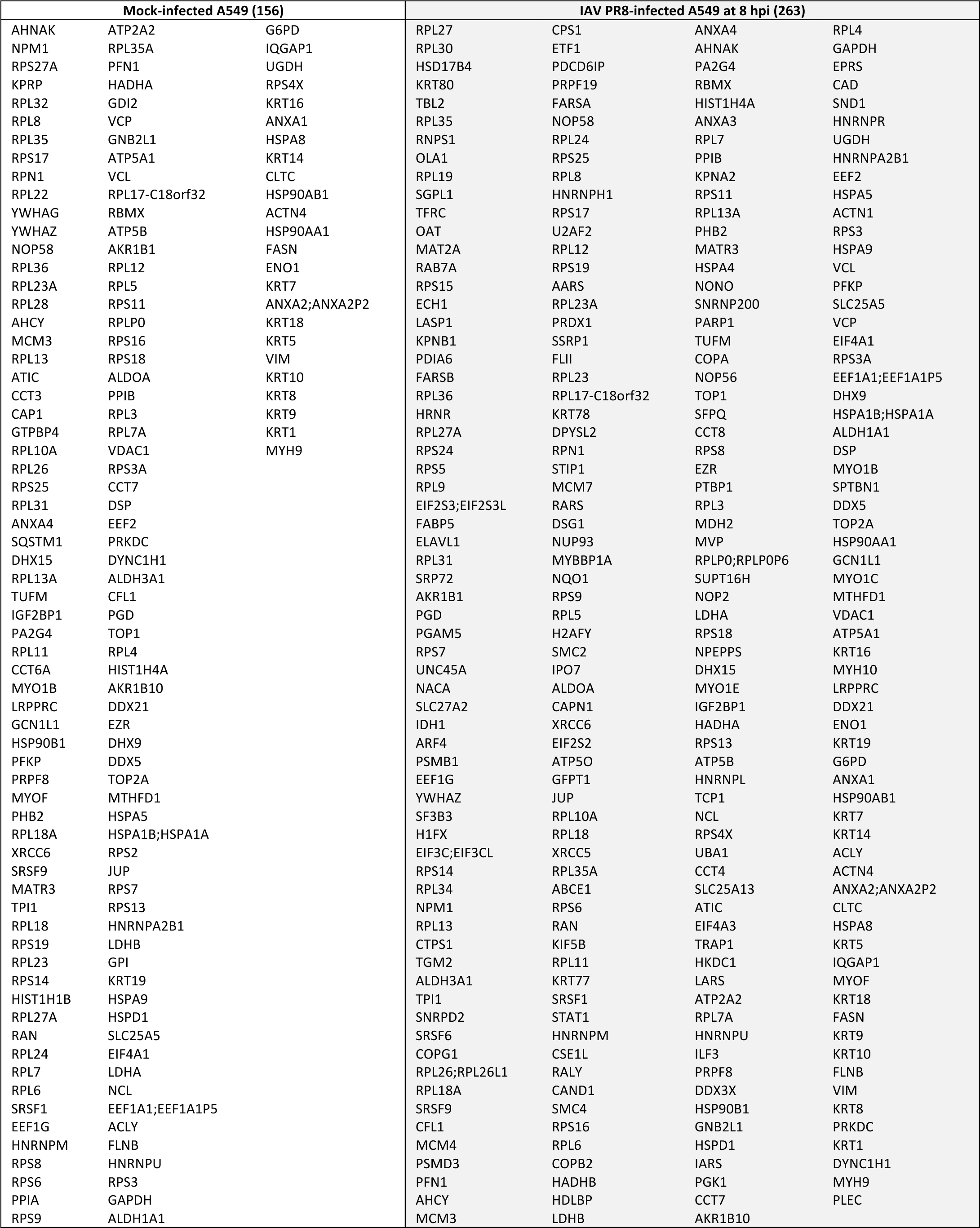
| Lists of host proteins identified in mock- and IAV PR8-infected A549 cells at 8 hpi (n=3) Proteins were considered when presenting at least 2 peptides and 2 unique peptides, in at least two of the experiments.

**Supplementary Table 3.**
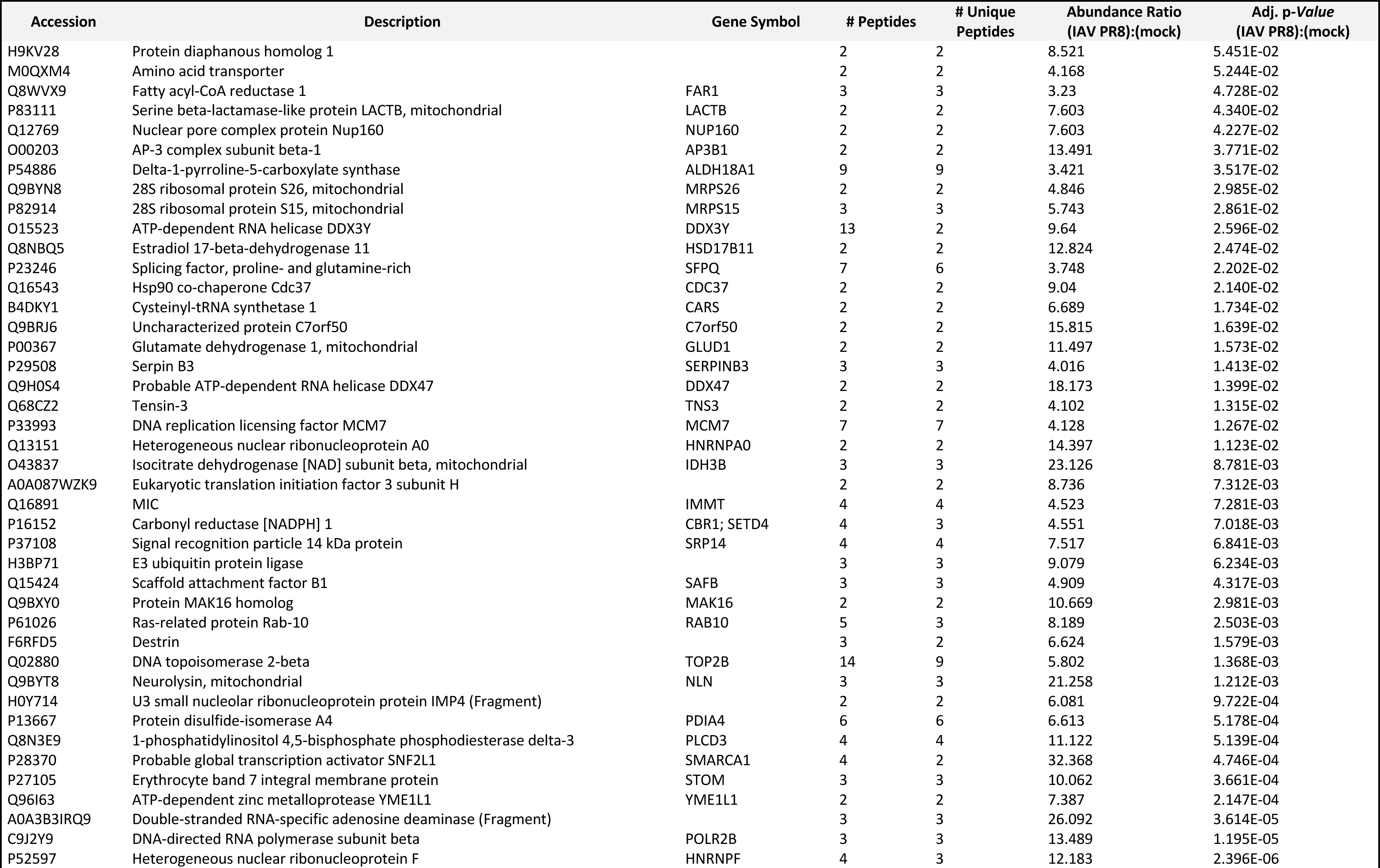
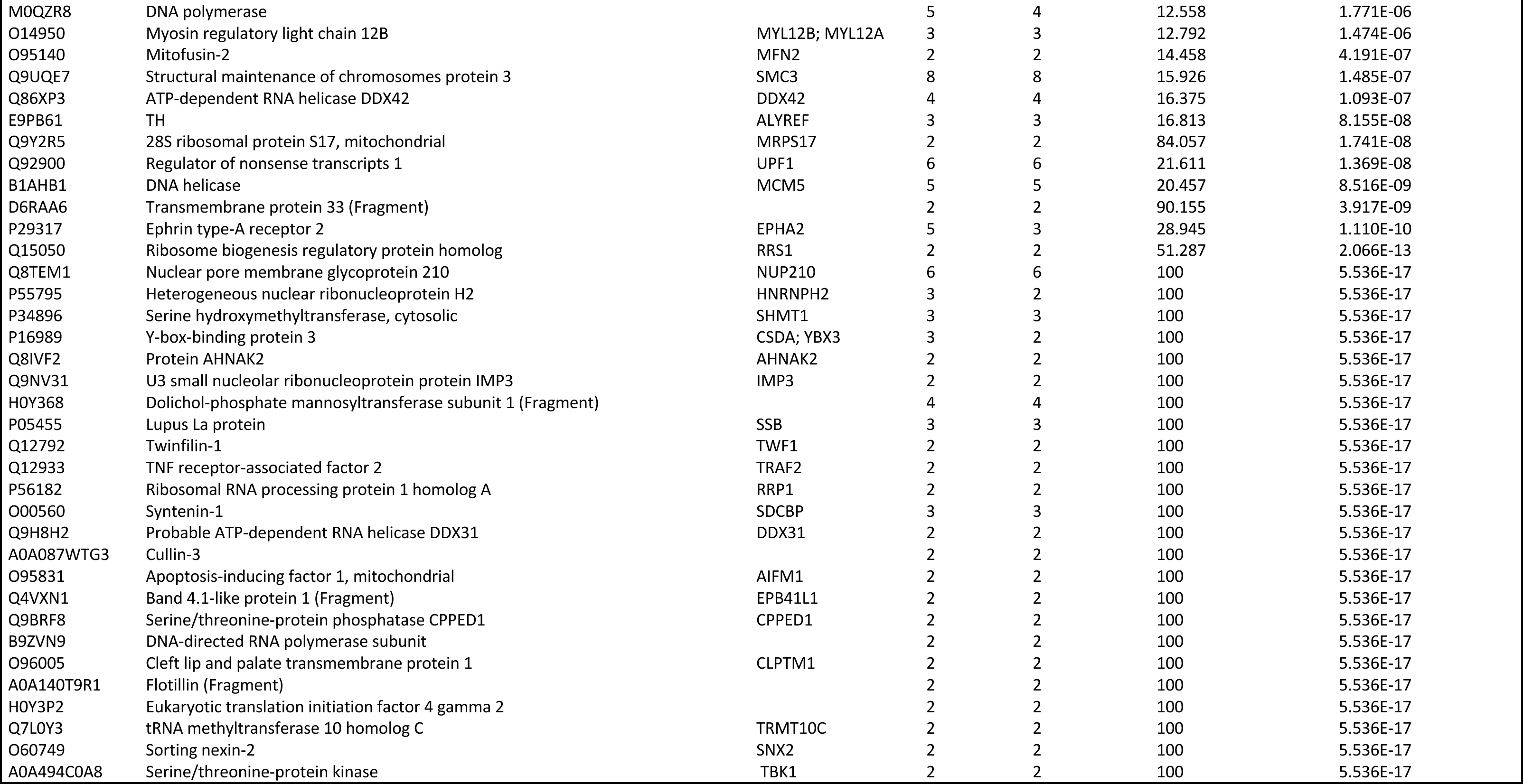
| Highly enriched host proteins in the insoluble protein fractions of influenza A virus-infected A549 cells at 8 hpi (n=3) Proteins were considered to be enriched in the insoluble protein fractions in infected samples when having the number of peptides and unique peptides ≥ 2, an abundance ratio (IAV PR8) / (mock) ≥ 1.5, and an abundance ratio adjusted p-value (IAV PR8) / (mock) ≤ 0.05.

**Supplementary Table 4.**
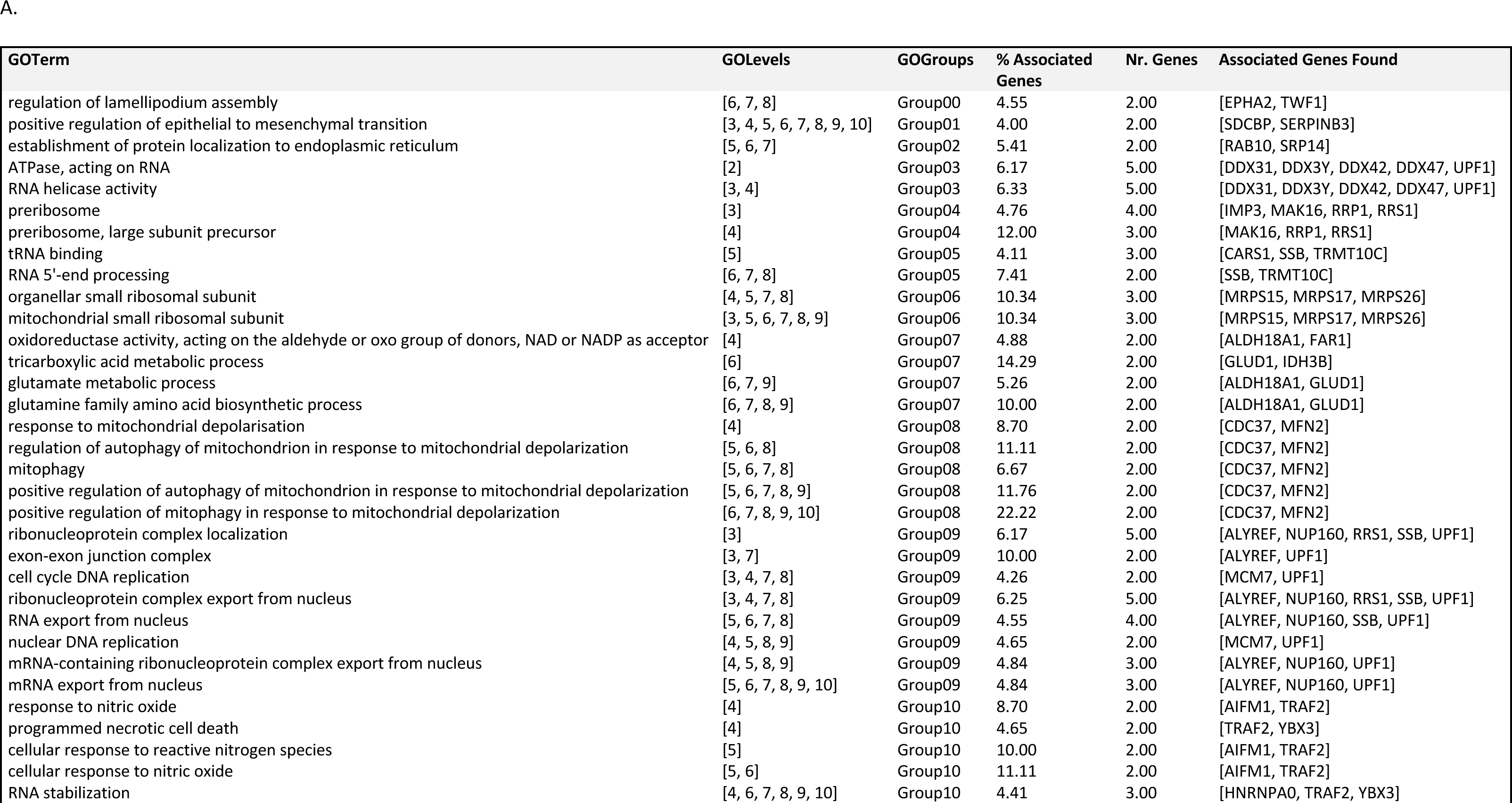
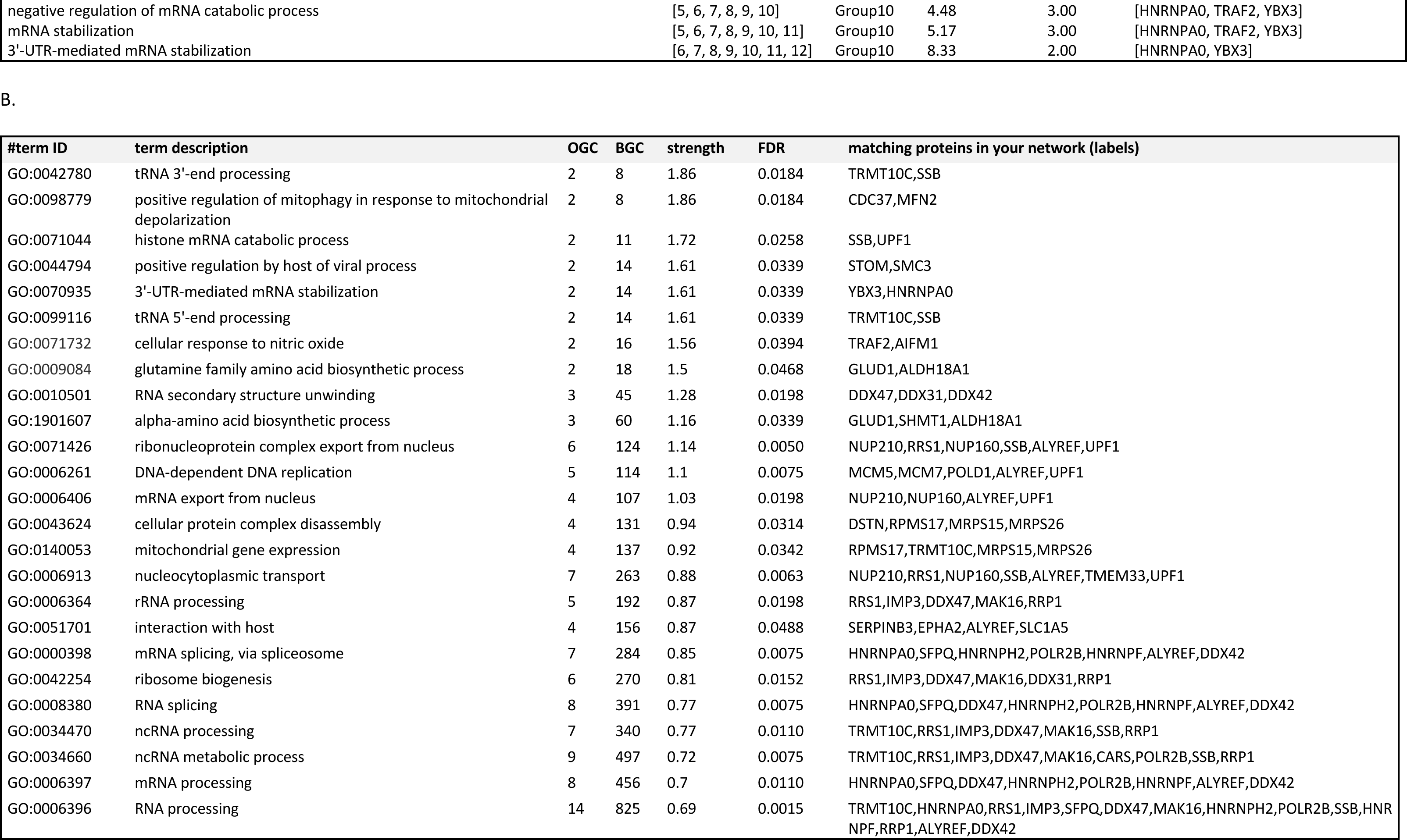
| Gene ontology analysis of proteins enriched in IAV PR8-infected samples in comparison to mock-infected A549 cells. (A) Analysis performed using Cytoscape ClueGo, considering biological processes, cellular components, and molecular function. Terms and groups presented have correspondent *p-*value<0.05. (B) TOP25 terms identified by STRING analysis (https://string-db.org/) considering biological processes. Observed gene count (OGC) indicates how many proteins are annotated with a particular term and background gene count (BGC) refers to how many proteins have this term assigned (in our list and in the background). Strength indicates the log_10_(observed/expected) and how large the enrichment effect is. The false discovery rate (FDR) measures how significant the enrichment is, showing the *p-values* corrected for multiple testing within each category using the *Benjamini-Hochberg* procedure.

**Supplementary Table.**
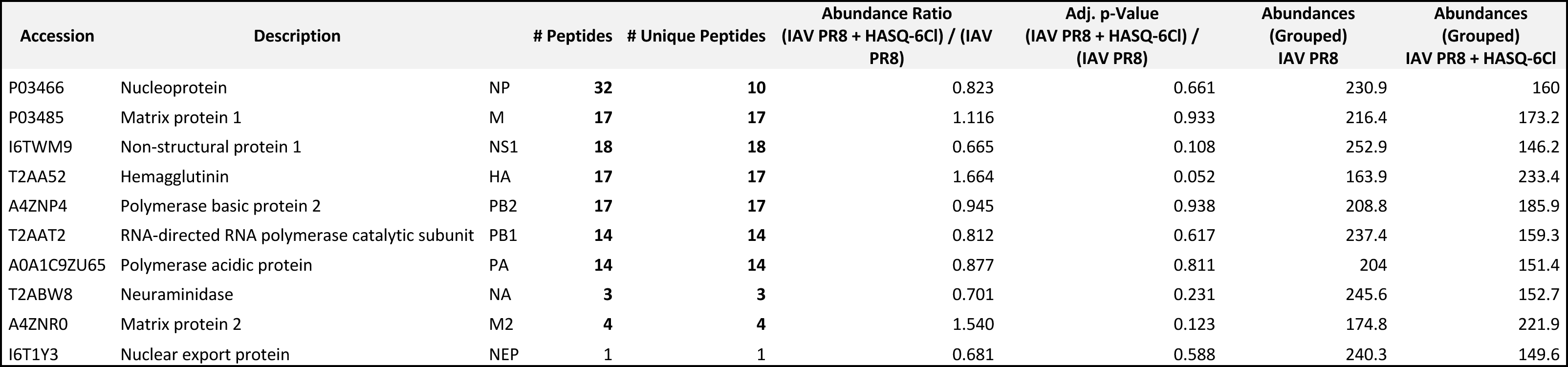
5 | Abundances of influenza virus (strain A/Puerto Rico/8/1934 H1N1) proteins in the insoluble protein fractions of infected A549 cells at 8 hpi in the presence or in the absence of HASQ-6Cl. Abundances (Grouped) from IAV PR8 and (IAV PR8 + HASQ-6Cl).

**Supplementary Table 6.**
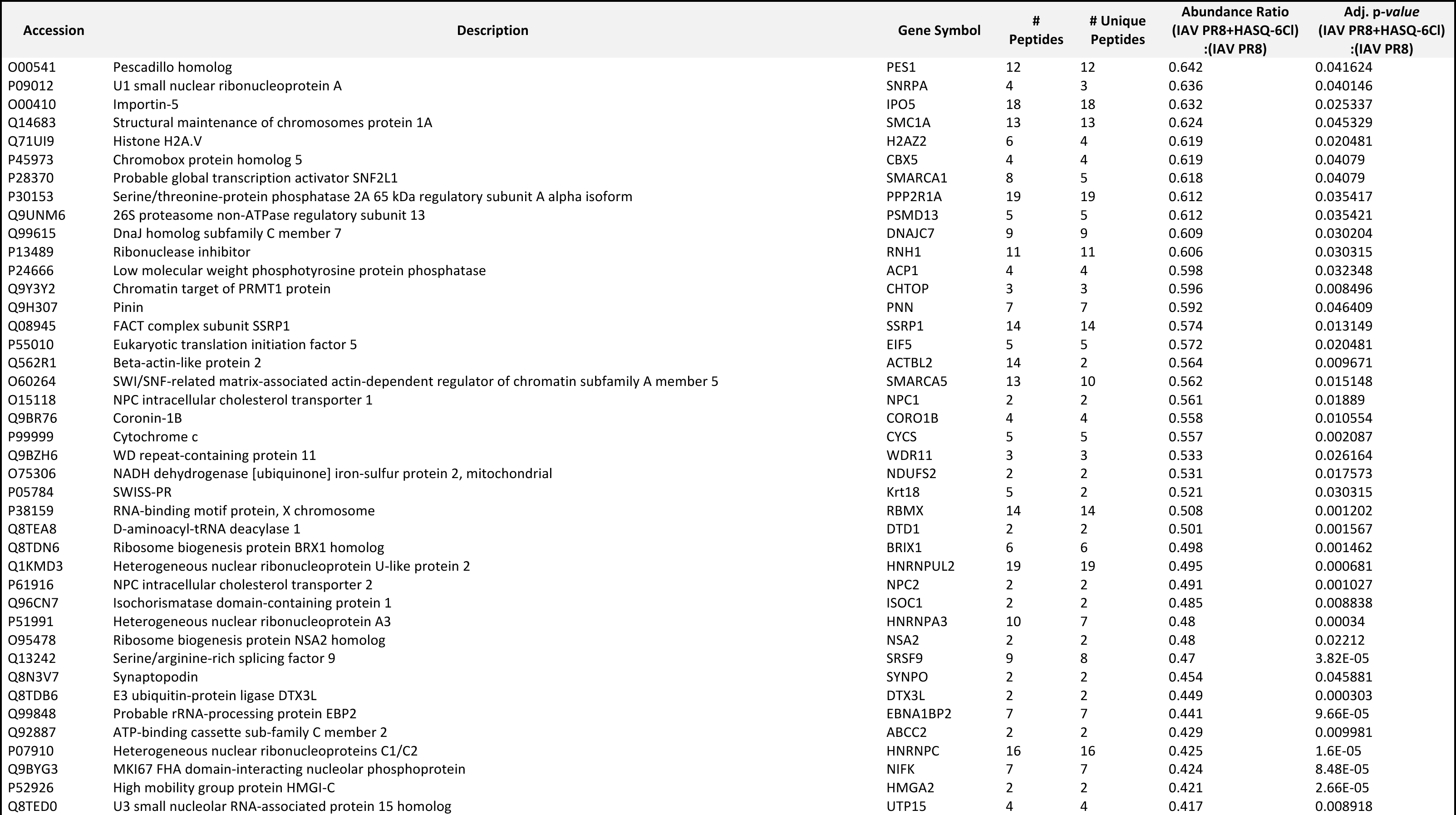
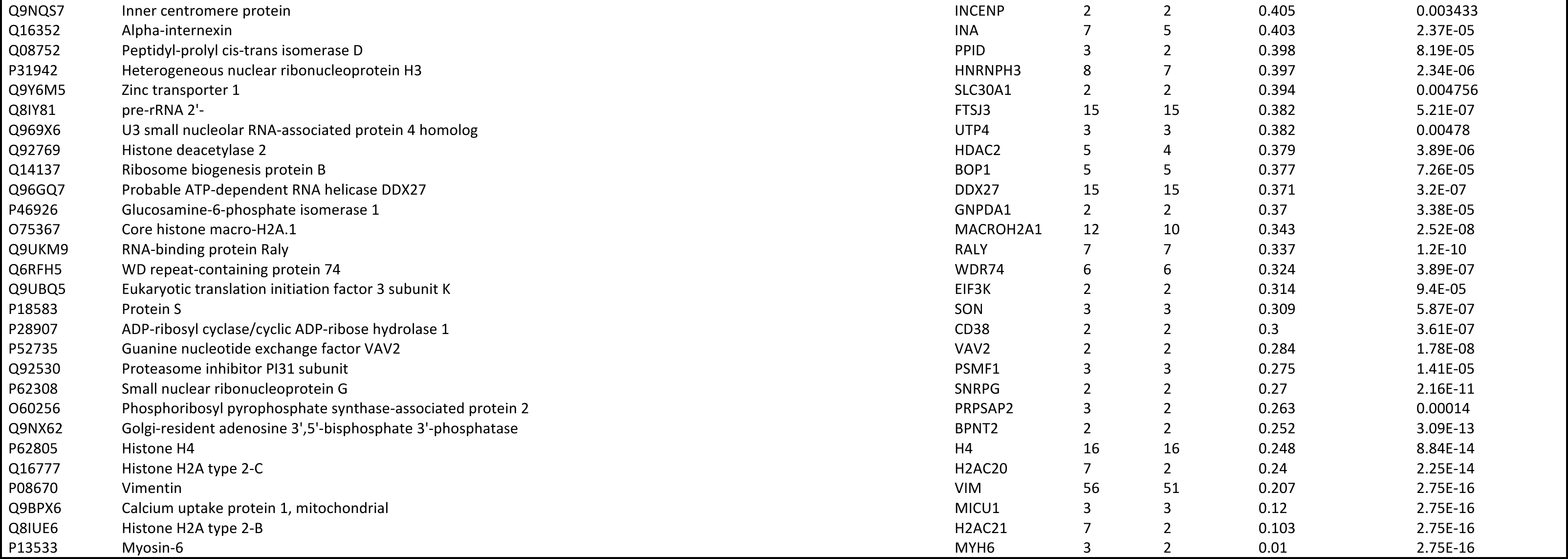
| Enriched host proteins in the insoluble protein fractions of influenza A virus-infected A549 cells at 8 hpi in comparison to IAV PR8-infected cells pre-treated with HASQ-6Cl (n=3) Proteins were considered to be enriched in the insoluble protein fractions in infected samples pre-treated with HASQ-6Cl when having the number of peptides and unique peptides ≥ 2, an abundance ratio (IAV PR8 + HASQ-6Cl) / (IAV PR8) ≤ 0.65, and an abundance ratio adjusted *p-value* (IAV PR8 + HASQ-6Cl) / (IAV PR8) ≤ 0.05.

**Supplementary Table 7.**
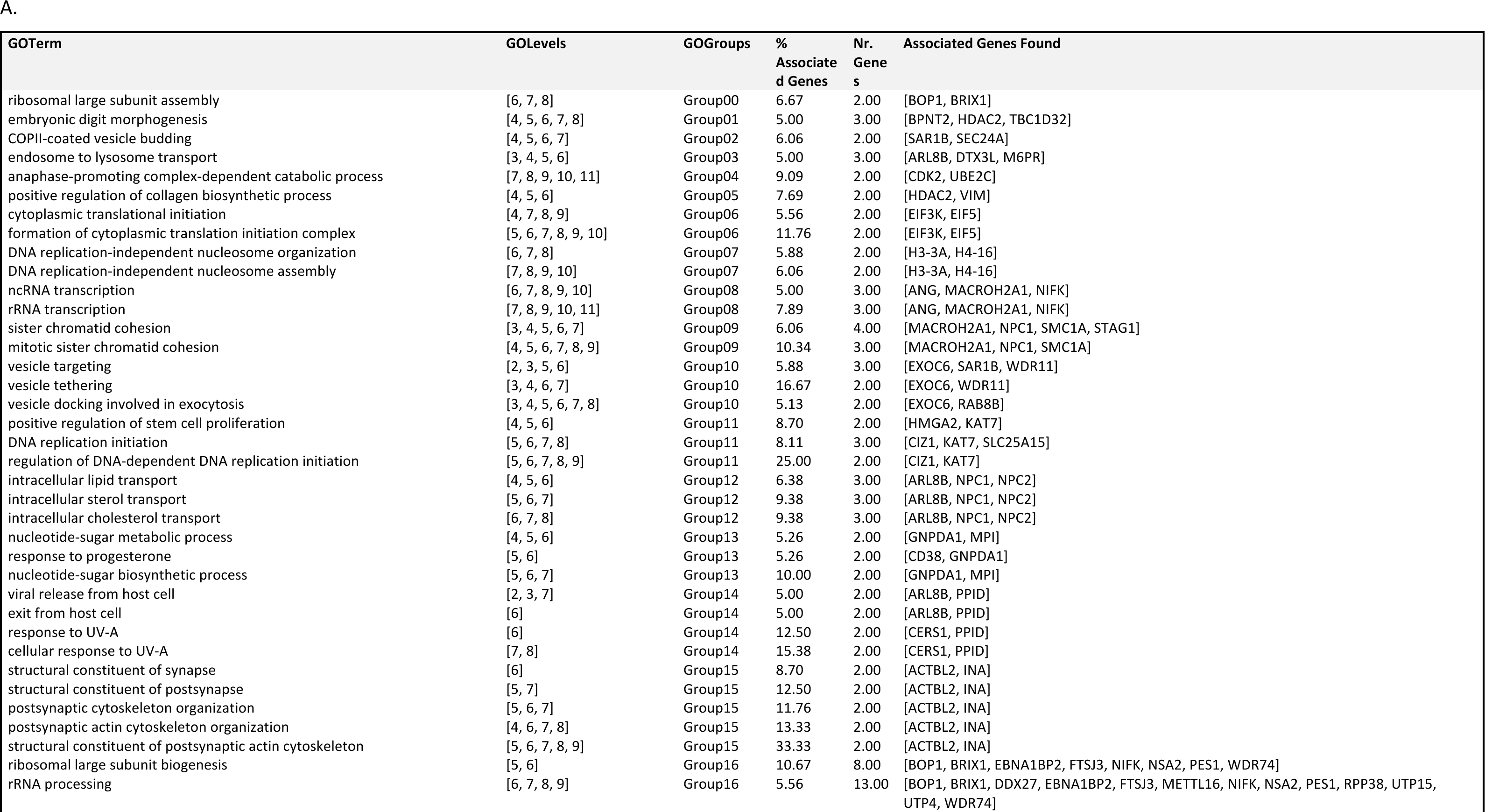
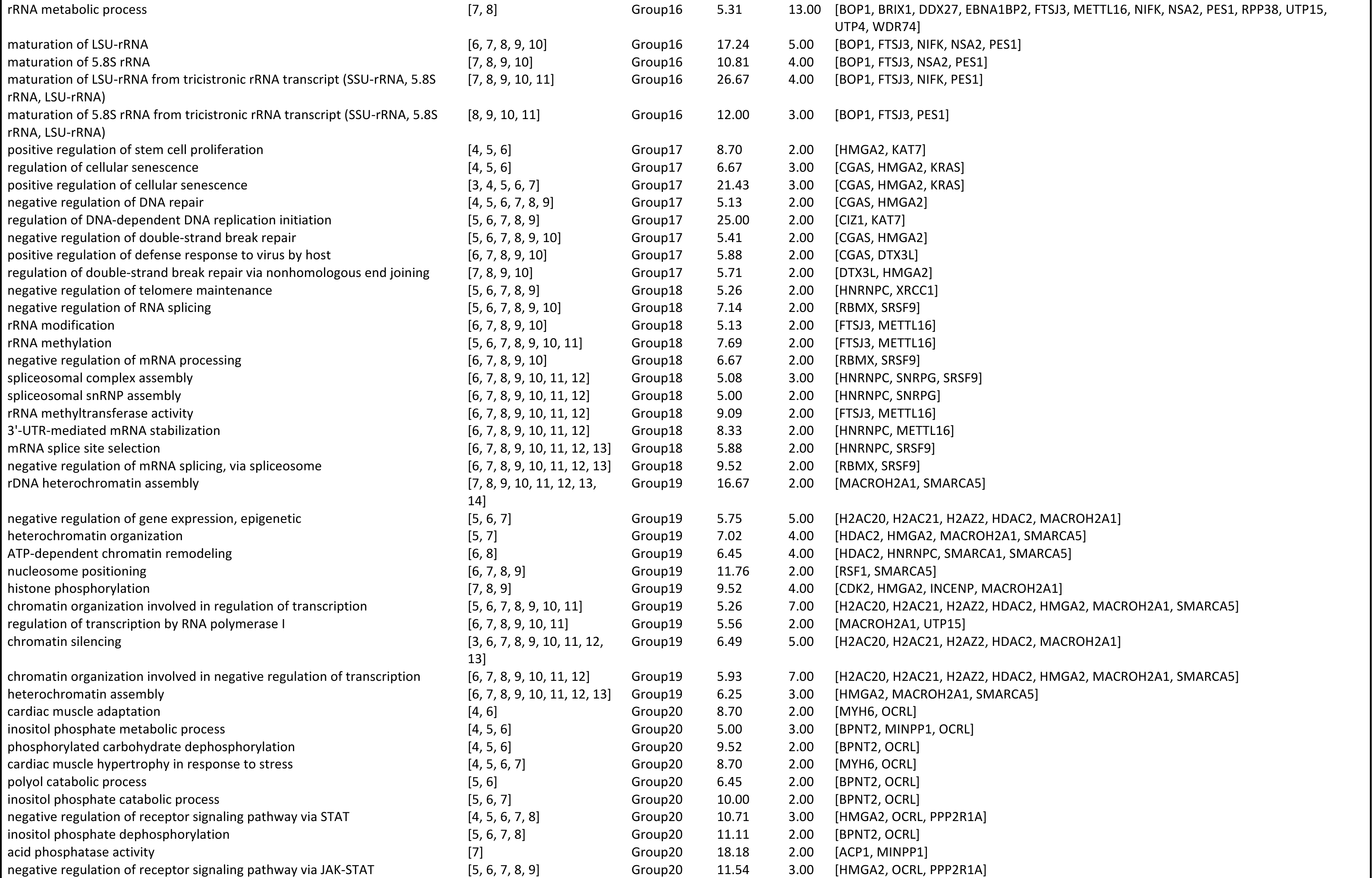
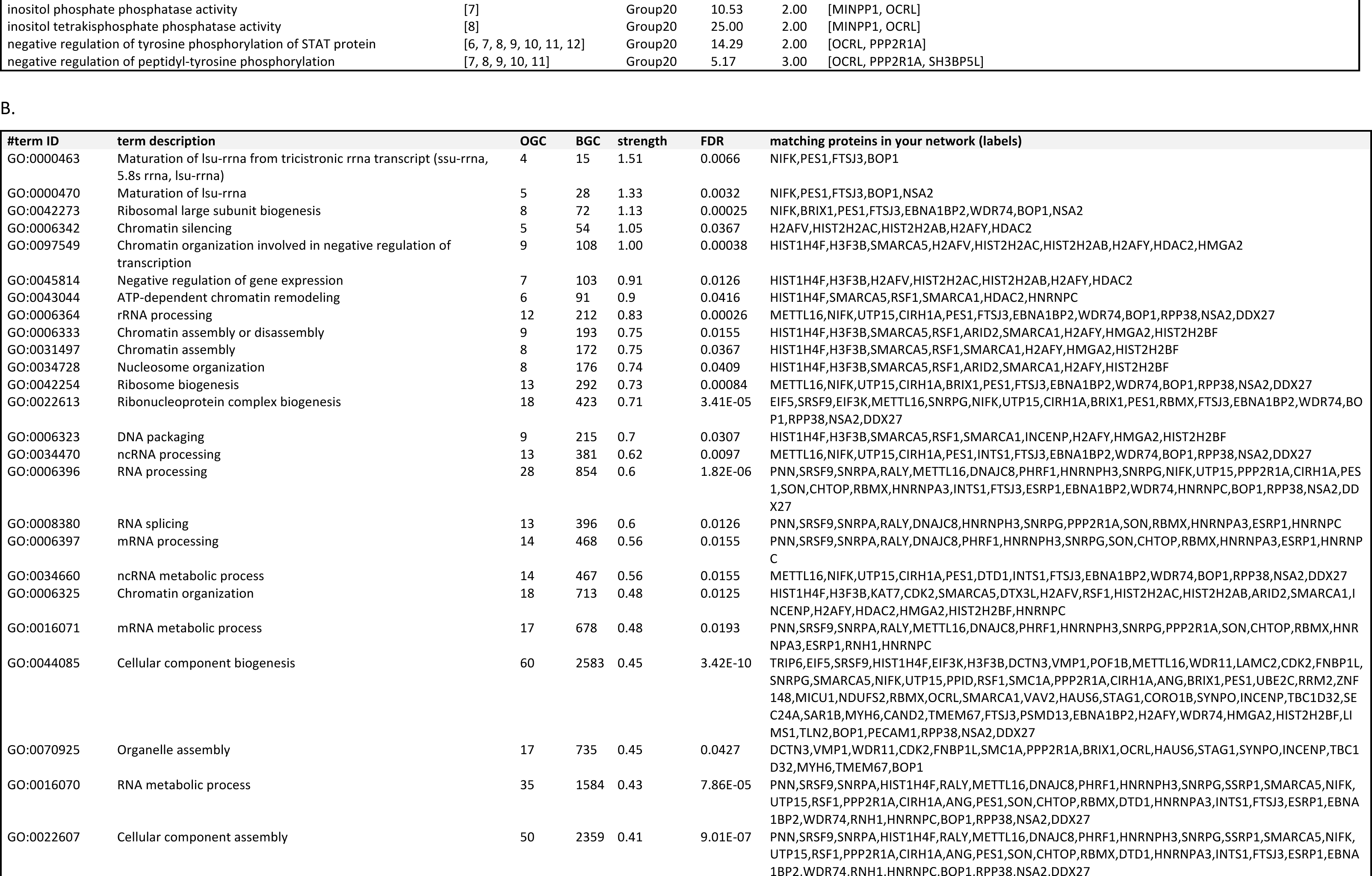
| Gene ontology analysis of proteins enriched in IAV PR8-infected samples in comparison to IAV PR8-infected cells pre-treated with HASQ-6Cl. ((A) Analysis performed using Cytoscape ClueGo, considering biological processes, cellular components, and molecular function. Terms and groups presented have correspondent *p-*value<0.05. (B) TOP25 terms identified by STRING analysis (https://string-db.org/) considering biological processes. Observed gene count (OGC) indicates how many proteins are annotated with a particular term and background gene count (BGC) refers to how many proteins have this term assigned (in our list and in the background). Strength indicates the log_10_(observed/expected) and how large the enrichment effect is. The false discovery rate (FDR) measures how significant the enrichment is, showing the *p-values* corrected for multiple testing within each category using the *Benjamini-Hochberg* procedure.

## Notes

### Competing Interest Statement

The authors have declared no competing interest.

